# Clinically oriented prediction of patient response to targeted and immunotherapies from the tumor transcriptome

**DOI:** 10.1101/2022.02.27.481627

**Authors:** Gal Dinstag, Eldad D. Shulman, Efrat Elis, Doreen S. Ben-Zvi, Omer Tirosh, Eden Maimon, Isaac Meilijson, Emmanuel Elalouf, Boris Temkin, Philipp Vitkovsky, Eyal Schiff, Danh-Tai Hoang, Sanju Sinha, Nishanth Ulhas Nair, Joo Sang Lee, Alejandro A. Schäffer, Ze’ev Ronai, Dejan Juric, Andrea B. Apolo, William L. Dahut, Stanley Lipkowitz, Raanan Berger, Razelle Kurzrock, Antonios Papanicolau-Sengos, Fatima Karzai, Mark R. Gilbert, Kenneth Aldape, Padma S. Rajagopal, Tuvik Beker, Eytan Ruppin, Ranit Aharonov

## Abstract

**Background:** Precision oncology is gradually advancing into mainstream clinical practice, demonstrating significant survival benefits. However, eligibility and response rates remain limited in many cases, calling for better predictive biomarkers.

**Methods:** We present ENLIGHT, a transcriptomics-based computational approach that identifies clinically relevant genetic interactions and uses them to predict a patient’s response to a variety of therapies in multiple cancer types, without training on previous treatment response data. We study ENLIGHT in two translationally oriented scenarios: *Personalized Oncology (PO)*, aimed at prioritizing treatments for a single patient, and *Clinical Trial Design (CTD*), selecting the most likely responders in a patient cohort.

**Findings:** Evaluating ENLIGHT’s performance on 21 blinded clinical trial datasets in the PO setting, we show that it can effectively predict a patient’s treatment response across multiple therapies and cancer types. Its prediction accuracy is better than previously published transcriptomics-based signatures and is comparable to that of supervised predictors developed for specific indications and drugs. In combination with the IFN-γsignature, ENLIGHT achieves an odds ratio larger than 4 in predicting response to immune checkpoint therapy. In the CTD scenario, ENLIGHT can potentially enhance clinical trial success for immunotherapies and other monoclonal antibodies by excluding non-responders, while overall achieving more than 90% of the response rate attainable under an optimal exclusion strategy.

**Conclusion:** ENLIGHT demonstrably enhances the ability to predict therapeutic response across multiple cancer types from the bulk tumor transcriptome.

**Funding:** This research was supported in part by the Intramural Research Program, NIH and by the Israeli Innovation Authority.

## Introduction

The current paradigm of precision oncology (PO), rooted in the 1990s development of trastuzumab and imatinib, focuses on matching individual targets to molecularly derived therapies ^1,2^. In the past 20 years, cancer therapeutics, driven by treatments developed for specific oncogenes and by the advent of immunotherapy, has overwhelmingly dominated regulatory drug approvals ^3^. For patients with a qualifying biomarker and otherwise limited treatment options, this paradigm can demonstrate improvement in clinical outcomes ^4–7^, as in the I-PREDICT study ^8^, which uses DNA biomarkers to identify novel combination therapies for patients. However, large studies such as NCI-MATCH demonstrate the limitations of the variant-centered biomarker approach, with less than 20% of patients ultimately assigned to single-therapy treatment arms ^9^. Response rates in the setting of targeted therapy also show broad ranges, between 25-75% of patients ^10^. Although qualification for immunotherapy varies by cancer type, often only 20-40% of patients respond to treatment ^11,12^.

One strategy to broaden which patients qualify for treatment and to improve response rates is to leverage omics data beyond genomics, with the main focus of current research being on transcriptomics. Clinical programs such as University of Michigan’s MIONCOSEQ have sought to integrate DNA and RNA-based sequencing to validate and support existing biomarkers ^13,14^. WINTHER was a prospective umbrella trial in which patients received personalized monotherapy or combination therapy based on genomic or transcriptomic data ^15^. This was the first clinical trial to examine the utility of RNA sequencing (RNA-seq) in a prospective clinical setting by targeting highly expressed cancer-associated genes, demonstrating that the tumor transcriptome could be used to provide complementary clinical information to DNA.

Other attempts to identify predictors of response from the transcriptome have only been studied in limited contexts to date ^16–19^. Notable examples include Sammut et al. ^*20*^ who demonstrated the ability of RNA-seq to improve prediction of pathologic complete response in early breast cancer (a space where RNA-seq has already demonstrated clinical utility ^21–23^) beyond DNA-seq, pathology imaging and clinical data; Jiang et al. ^*16*^ and Cui et al. ^*24*^ who built predictors for response to immune checkpoint blockade based on the tumor’s immune microenvironment, reflected by RNA data; and the OncoTarget / OncoTreat program at Columbia University that builds on using RNA-seq alone by mapping protein interaction networks derived from the transcriptome to prioritize cancer drivers for treatment and was evaluated in pancreatic neuroendocrine tumors ^25^. These important and novel approaches have, however, been limited to highly specific clinical contexts and treatments and each has different data requirements. Using such approaches in a scenario where multiple treatment options per patient exist is therefore a very challenging task ^12,26–28^.

In a recent effort to overcome these limitations and develop a uniform systematic approach for stratifying patients to multiple therapies based on the tumor transcriptome, Lee et al. ^*29*^ have demonstrated that synthetic lethality (SL) and synthetic rescue (SR) interactions can be leveraged to predict treatment response via the transcriptome. An SL interaction between two genes means that the simultaneous inactivation of both genes reduces the viability of the cell while the individual inactivation of each does not ^30,31^. An SR interaction is one in which the inactivation of one gene reduces cell viability, but an alteration of another gene’s activity, termed the *rescuer*, restores (rescues) viability ^32,33^. Their framework, SELECT, mines large-scale in-vitro and TCGA patient data to computationally infer putative pairs of Genetic Interactions and uses these interactions to predict drug response and prioritize a variety of treatments. It showed that tumor expression could be analyzed systematically to predict patients’ response to a broad range of targeted therapies and immunotherapies in multiple cancer types with high accuracy. However, SELECT has a few notable limitations, which we have set to remedy.

To this end, here we substantially extend the work of Lee et al. and present *ENLIGHT*, a transcriptomics-based precision oncology pipeline based on the GI-networks approach. First, our work has been motivated by a survey we have conducted, where we evaluated the ability of SELECT to identify GI networks for the genes that are inhibited by 105 FDA approved targeted and immunotherapies, finding that it could build such GI networks for only 67% (70/105) of them, thus limiting its possible use in clinical settings. Addressing this challenge with a new GI-based scoring algorithm that considers different types of GIs concomitantly, ENLIGHT successfully extends the scope of SELECT to *all* these 105 therapies. Second, SELECT had focused on maximizing the area under the ROC curve (AUC) – a measure commonly used in data science but not directly applicable in clinical settings. In contrast, ENLIGHT’s design and parameter optimization is aimed at maximizing its performance on three key translational tasks: (i) *Drug coverage*– determining the range of drugs for which predictions can be obtained and extending the prior range of SELECT; (ii) *Personalized oncology*– evaluating and prioritizing multiple candidate treatment options for individual patients. Since a variety of treatment options are considered in this scenario, a good test should primarily have a high Positive Predictive Value (*PPV*, also known as *precision*) taking priority over *sensitivity (recall*), which can be moderate; (iii) And finally, improving *Clinical Trial Design (CTD*) by excluding patients that are unlikely to respond to the treatment by employing a test that has a high Negative Predictive Value (*NPV*).

To evaluate ENLIGHT’s performance, we analyzed 21 new cohorts not analyzed in the SELECT study, overall spanning 697 patients receiving 15 different drugs in 11 cancer indications, and importantly, kept them blinded for evaluation purposes. We find that while SELECT is reassuringly predictive on the subset of those datasets for which it could produce results, ENLIGHT still performs better when evaluated using clinically relevant performance measures. We show that (i) ENLIGHT produces predictions for all the drugs used in these cohorts; (ii) High *ENLIGHT-Matching Scores (EMS*) are associated with better response (odds ratio (OR) = 2.59; 95% confidence interval [1.85, 3.55]; *p*=3.41*e*-8). (iii) ENLIGHT can be used to successfully exclude patients from clinical trials, achieving overall more than 90% of the response rate attainable under an optimal exclusion strategy for both immunotherapies and targeted monoclonal antibody (mAb) treatments. Finally, we find that *ENLIGHT performs as well as supervised tests developed for specific indications and drugs* and better than other known transcriptomics-based predictive and prognostic signatures. These results showcase the effectiveness of ENLIGHT as a tool to improve translational oncology research.

## Results

### Tuning and Evaluation Datasets

To tune the parameters for ENLIGHT, including the GI network size and a *decision threshold* on the EMS, that is used for predicting a favorable response (see below), we selected eight *tuning sets –* six datasets ^34–39^ already analyzed in Lee et al. ^*29*^ along with two other datasets ^40,41^ from the public domain. These were selected as they span a range of different treatments, therapeutic classes, response rates and sample sizes, reflecting diverse real-world data, covering five targeted therapies and one immune checkpoint blockade (ICB) therapy.

To study the value of ENLIGHT on real world data, which was not previously analyzed by either SELECT or ENLIGHT, we surveyed the public domain (GEO ^42^, ArrayExpress ^43^, CTRDB ^44^ and the broader literature) for available datasets of patients receiving targeted or immunotherapies, containing both pre-treatment transcriptomics and response information. In addition, we publish here a new dataset of breast cancer patients who received *Alpelisib* or *Ribociclib* that was obtained as part of a collaboration with Massachusetts General Hospital (MGH). Overall, we identified, obtained and set aside 21 datasets ^20,24,34,39,45–61^–*the evaluation sets*– as unseen data for evaluation. See **Table S1** and **STAR Methods** for full details.

### Overview of the ENLIGHT Pipeline

ENLIGHT’s drug response prediction pipeline generally follows the flow of SELECT ^29^ which comprises two steps (**Figure 1a**): (i) Given a drug, the *inference engine* identifies the clinically relevant GI network of the drug’s target genes. As was done in SELECT, we start with a list of initial candidate SL/SR pairs by analyzing cancer cell line dependencies with RNAi, CRISPR/Cas9, or pharmacological inhibition, based on the principle that SL/SR interactions should decrease/increase tumor cell viability, respectively, when *activated* (e.g., in the SL case, when both genes are downregulated). Among these candidate pairs, we select those that are more likely to be clinically relevant by analyzing a database of tumor samples with associated transcriptomics and survival data, requiring a significant association between the joint inactivation of a putative SL gene pair and better patient survival, and analogously for SR interactions. Finally, among the candidate pairs that remain after these steps, we select those pairs that are supported by a phylogenetic profiling analysis (see **STAR Methods** for more details). When considering combination therapies, ENLIGHT computes a GI network based on the union of all the drug targets. (ii) The drug-specific GI network is then used to *predict* a given patient’s response to the drug based on the gene expression profile of the patient’s tumor. The ENLIGHT Matching Score (EMS), which evaluates the match between patient and treatment, is based on the overall activation state of the genes in the drug target’s GI network, reflecting the notion that a tumor would be more susceptible to a drug that induces more active SL interactions and fewer active SR interactions. An important prerequisite for a drug to be analyzed by this approach is that the gene targets are well defined. Thus, in this work we focus on targeted therapies and ICBs.

**Figure 1:**
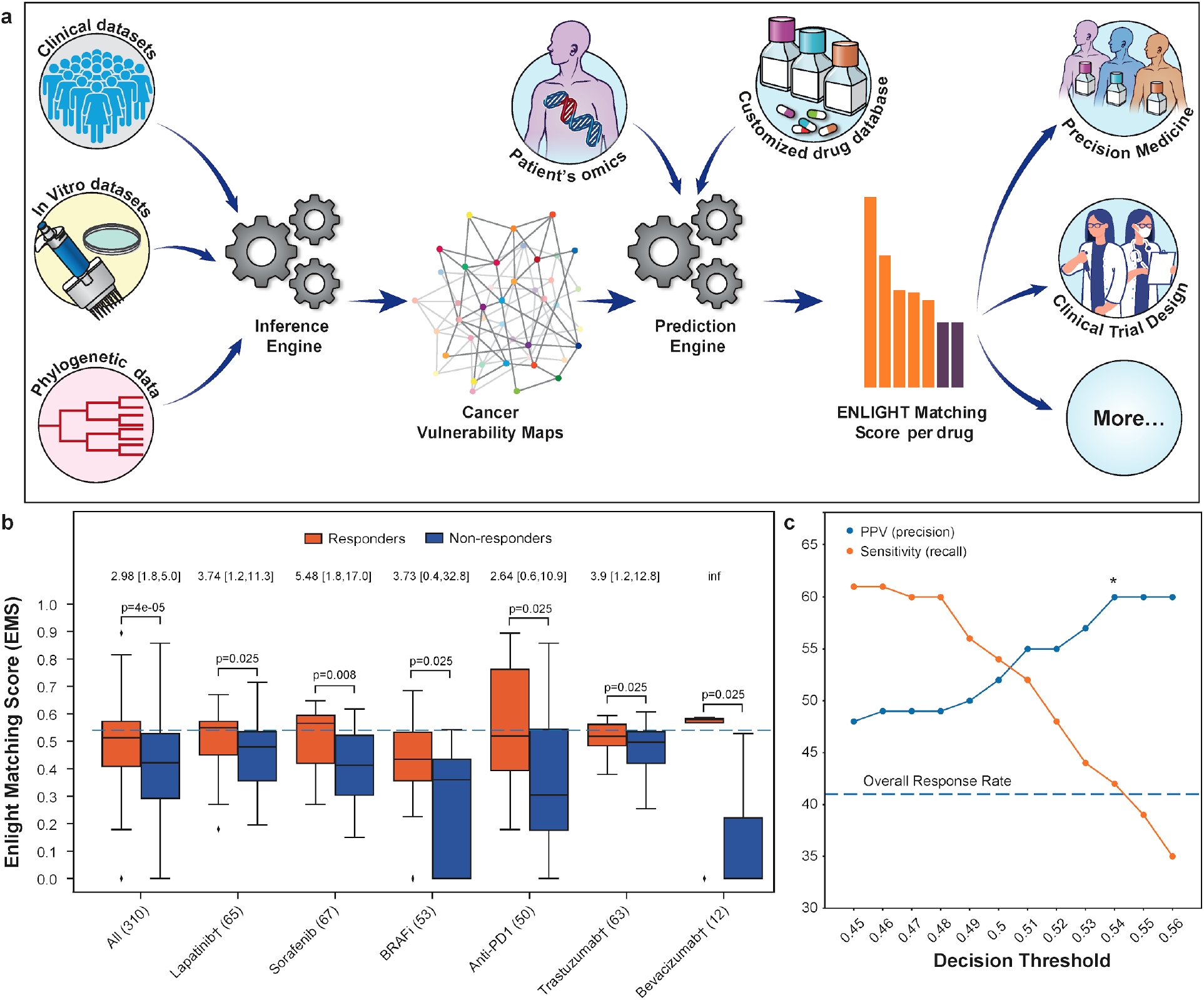
The ENLIGHT Pipeline and its tuning for Personalized Oncology. **(a)** ENLIGHT’s pipeline starts by inferring the GI network of a given drug. The GI network and omics data from patients are then used to predict treatment response. **(b)** ENLIGHT matching scores in responders (orange) are higher than in non-responders (blue), across the tuning cohorts. The 3 BRAFi cohorts are very small (N = 16, 17 and 20), so their results are presented in aggregate. The ‘All’ column presents the results for all patients aggregated together. p-values were calculated using a one-sided Mann-Whitney test. The horizontal line marks the EMS decision threshold. The odds ratio of ENLIGHT-matched cases appears on top of each pair of bars, with brackets indicating the 95% confidence interval. The number of patients in each cohort is provided in parentheses. **(c)** The PPV (precision) and sensitivity (recall) on the aggregate tuning cohort (y axis) as a function of the decision threshold (x axis).

In developing ENLIGHT, we have extended and improved this basic framework, by introducing the following adaptations (see **Supplementary Note 1** for more details): **(i)** ENLIGHT’s GI networks include both SL and SR interactions that are concomitantly inferred for each drug, thereby considerably increasing drug coverage relative to SELECT, which utilizes only one type of interaction per network (for immunotherapy, where SELECT had no coverage limitation, we use the same SR interactions as in Lee et al., see Figure 5a in ^29^ for details). **(ii)** To improve the inference engine, we follow the idea presented in Lee et al.^*62*^ and Sahu et al. ^*63*^ and add a depletion test (not present in SELECT), requiring that the joint inactivation of a candidate SL pair is under-represented in tumors (and analogously for SR partners). **(iii)** SELECT has used a Cox proportional hazard test on categorized expression data to identify candidate SL/SR pairs that confer favorable/unfavorable patient survival when the interaction is active. To increase robustness and statistical power, ENLIGHT applies a fully parametric test, based on an exponential survival model, on continuous expression data. **(iv)** ENLIGHT GI networks are considerably larger than those of SELECT to reduce score variation across drugs and indications. **(v)** To improve the prediction engine, for immunotherapy and other mAbs, which are highly target-specific, the EMS incorporates the expression of the target as an additional score component. The ENLIGHT pipeline is described in detail in **STAR Methods**, where we also describe a web service enabling researchers to apply ENLIGHT to RNA expression data.

We emphasize that the GI networks are inferred without using any treatment response data. Notably, the parameters of ENLIGHT, namely the network size and thresholds for the inference tests, are optimized on the tuning datasets, but their values are then held fixed to predict all treatment outcomes across all 21 evaluation sets. These conservative procedures markedly mitigate the risk of deriving overfitted predictors that would not predict response in new datasets.

### Setting the ENLIGHT Decision Threshold for Personalized Oncology Based on the Tuning Cohorts

To define an ENLIGHT-based test for the *personalized oncology* scenario, we define a *uniform decision threshold* on the EMS, above which the probability of a patient to respond is high. The chosen decision threshold should maximize the proportion of true responders among those predicted to respond (the PPV), while the proportion of true responders identified by the test (termed its sensitivity, or recall) does not have to be very high since multiple treatment options are usually considered. In other words, in a real-life setting a physician needs to choose one of multiple treatment options, hence our objective is to maximize the PPV of the recommendations given that at least one match would be found for, ideally, each patient. Importantly, the decision threshold, set on the tuning cohorts, is fixed, and is used as is in evaluating ENLIGHT’s performance on all cohorts and treatments. The EMS distributions in the tuning cohorts show that the EMS are significantly higher among responders compared to non-responders (**Figure 1b**), testifying to its discriminatory power. **Figure 1c** shows that a decision threshold of **0.54**, which will be used henceforth, maximizes the PPV while still maintaining a reasonable sensitivity on the tuning cohort (for the dataset-specific PPV and sensitivity curves see **Figures S1** and **S2**). ENLIGHT’s performance is evaluated using the odds ratio (OR) for response, which denotes the odds to respond when the treatment is recommended (*ENLIGHT-matched*, i.e., EMS above the decision threshold), divided by the odds when it’s not. As shown in **Figure 1b**, ENLIGHT obtains an aggregate OR of 2.98 on the tuning cohorts (aggregation based on individual patients; 95% confidence interval [1.8, 5]; *p*=4e-05).

### ENLIGHT Successfully Predicts Patients Treatment Response in 21 Different Test Datasets

We next turned to evaluate how ENLIGHT performs in identifying the true responders in the 21 unseen patient cohorts that we have collected, spanning ICBs, mAbs, and targeted small molecules. Notably, the response data for all evaluation cohorts was unblinded only after finalizing the ENLIGHT pipeline including fixing the decision threshold and calculating EMS for all patients. **Figure 2a** shows that ENLIGHT-matched treatments are associated with better patient response (OR > 1) in all cohorts except for two (*Sorafenib_2_* and one ICB cohort), with an aggregate OR of 2.59 (95% confidence interval [CI], 1.89 to 3.55; *p*=3.41*e*-8, N=697). Correspondingly, **Figure 2b** shows that the overall PPV obtained for ENLIGHT-matched cases is markedly higher than the overall response rate (52% vs 38%, a 36.84% increase, *p*=3.30e-13, one sided proportion test, and see **Table S2** for a more detailed account). Interestingly, ENLIGHT is more accurate in immunotherapies and other mAbs vs. targeted small molecules, which aligns with its reliance on drugs that have accurate targets. More specifically, within the small molecules class, ENLIGHT is only less predictive in drugs with many targets (*Sorafenib*, a broad tyrosine kinase inhibitor and *MK2206*, a pan-*AKT* inhibitor). Notably, when a patient received a combination of targeted and chemotherapy agents (see cohorts marked with a cross), the EMS was calculated for the targeted agent alone, however, remarkably, the performance is still maintained.

**Figure 2:**
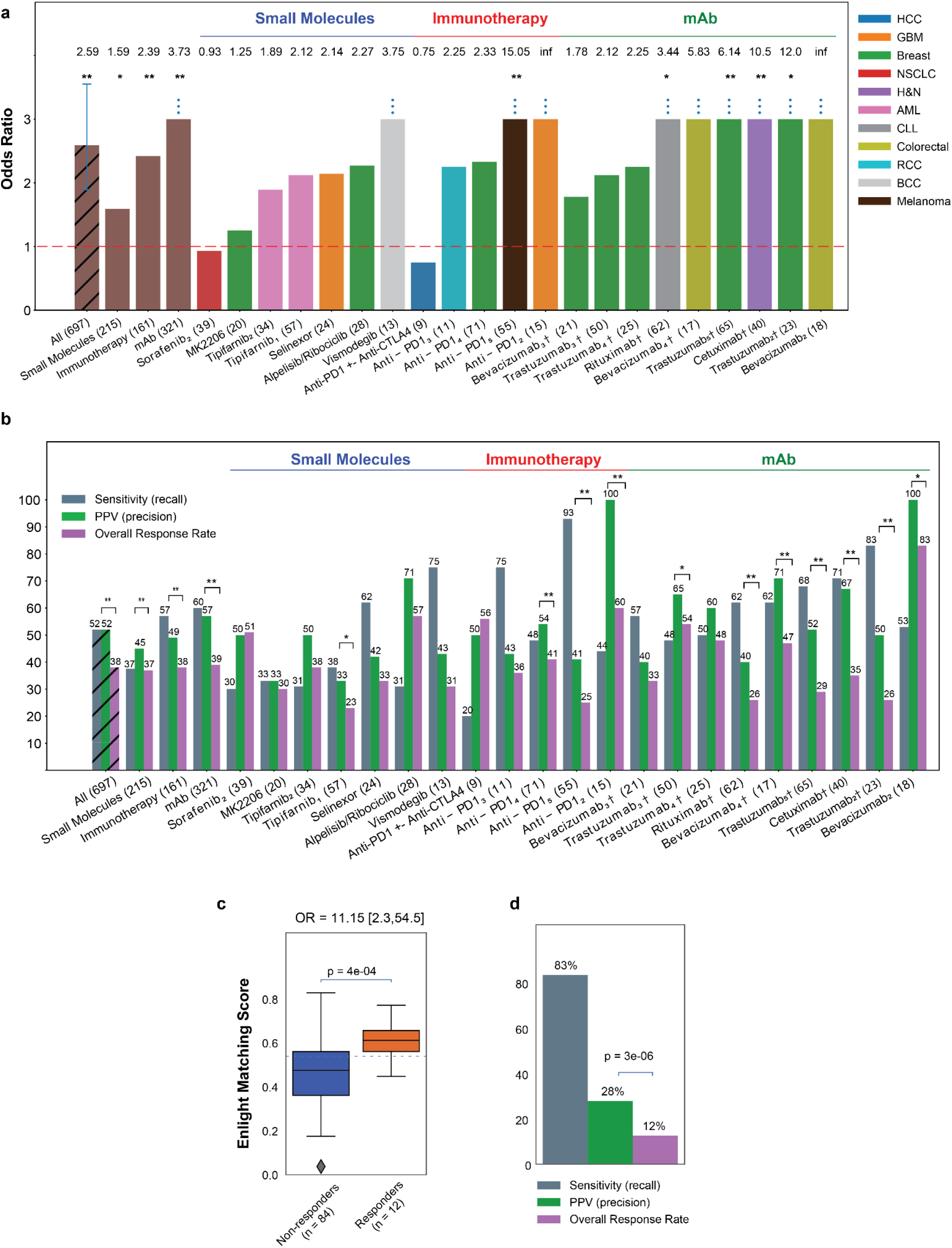
ENLIGHT’s ability to stratify patients for therapy. **(a)** The bar graphs show the odds ratio for response of ENLIGHT-matched cases in the 21 evaluation cohorts (OR values appear on top of each bar; all eight patients predicted to respond in the *Bevacizumab_2_cohort* responded to the treatment, resulting in an infinite OR), along with the OR for the aggregation of all cohorts and aggregation based on therapeutic-class. Sample sizes are denoted in parentheses. cohorts for which OR is significantly larger than 1 according to Fisher’s exact test are denoted with asterisks. “Anti-PD1” encompasses three different drugs (Nivolumab, Pembrolizumab and Durvalumab)-see details in **Table S1**. **(b)** Analogous to (a) but presenting the sensitivity and PPV of ENLIGHT-matched cases vs. the overall response rate for the evaluation cohorts and their aggregations. Significant differences between PPV and response rate according to the one-sided proportion test are denoted with asterisks. **(c)** In the WINTHER trial, responders (orange) have significantly higher EMS than non-responders (blue); p-value was calculated using a one-sided Mann-Whitney test. The horizontal line marks the decision threshold (0.54). **(d)** The sensitivity and PPV of ENLIGHT-matched cases vs. overall response rate in the WINTHER trial. [†: Patients in these cohorts received a combination of targeted and chemotherapy; *: p<0.1; **: p<0.05]

Additionally, we evaluated ENLIGHT as a personalized oncology tool in a multi-arm clinical trial setting, by analyzing data from the WINTHER trial, a large-scale prospective clinical trial that has incorporated genetic and transcriptomic data for cancer therapy decision making in adult patients with advanced solid tumors ^15^. ENLIGHT was able to provide predictions for all patients, except four (see **Supplementary Note 2**). The EMS of the responders were significantly higher than those of non-responders (*p*= 4e-04, **Figure 2c**). The odds ratio of ENLIGHT-matched treatments is 11.15 (*p*= 8e-04, **Figure 2c**), and the PPV is more than two times higher than the overall response rate (**Figure 2d**). Further analysis shows that responders had significantly higher EMS than non-responders also for the 24 patients treated with a combination of drugs (**Figure S3a**) and that ENLIGHT-matched treatments were associated with better response, without being hampered by the background of chemotherapy treatment (**Table S3**). **Figure S3b**depicts the landscape of different treatment alternatives with high EMS scores for each patient. We observe that 91/96 patients (94.8%) had at least one treatment with which they were ENLIGHT-matched, highlighting the potential coverage of ENLIGHT in real-world cases.

Except for tuning very few parameters on the tuning cohorts, as described above, ENLIGHT is essentially an *unsupervised prediction method* that relies on a series of biologically motivated statistical tests that underlie the inference of GIs and their utilization and is not trained on treatment outcome data. We next turned to compare its predictive power to that of transcriptomics-based biomarker signatures generated via supervised machine learning on specific training cohorts. To this end, we have studied four of the evaluation datasets, for which a supervised classifier was presented in the original publication and data enabling us to perform this comparison was provided (**Figure 3a**).In one of these datasets, Sammut et al. ^20^ used supervised learning to predict response to chemotherapy with or without *Trastuzumab* among HER2+ breast cancer patients. We focused on the subset of 56 patients who received *Trastuzumab* and for which the model’s scores were published. ENLIGHT achieved a slightly higher AUC than that reported in the original study, without ever training on these data, nor on any other data of response to *Trastuzumab*, and without using any clinical features. Similarly, Cui et al. ^24^ developed a biomarker for response to anti-PD1 in melanoma (see *Anti-PD1_5_* in **Table S1**) and Raponi et al. ^*54*^ and Raponi et al. ^55^ developed biomarkers for response to *Tipifarnib* in AML to which we compared ENLIGHT’s performance. Overall, In three of the four datasets, ENLIGHT’s predictive performance is comparable to that of the biomarkers developed using a supervised approach for a specific treatment and indication. However, in one of these datasets (“Raponi et al. Blood”), while ENLIGHT has a slightly higher PPV, its sensitivity is much lower than that achieved by the supervised method. Taken together, these results demonstrate the remarkable predictive power of ENLIGHT as a pan-cancer and pan-treatment response predictor, which quite successfully competes with supervised models specifically trained for narrow predictive scenarios.

**Figure 3:**
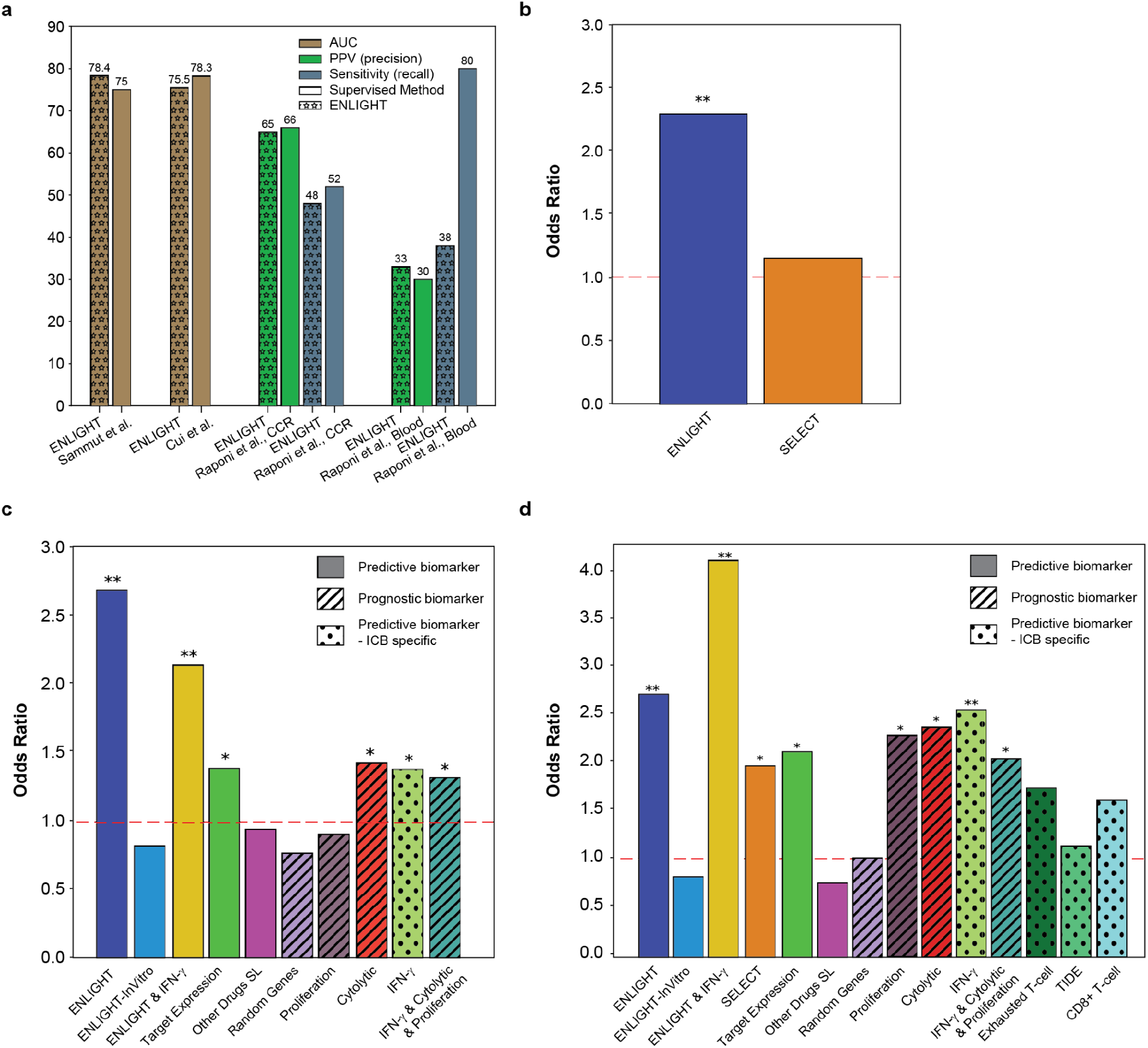
ENLIGHT’s predictive accuracy vs. other biomarkers. **(a)** Comparison of ENLIGHT’s performance to that of biomarkers developed using supervised methods. Comparison was performed on the performance measure presented in the respective publication. **(b)** Odds Ratio (OR, y-axis) of ENLIGHT (blue) and SELECT (orange) calculated for the 10 cohorts (N = 297) for which SELECT was able to generate a prediction. **(c)** OR (y-axis) of ENLIGHT (blue) and other transcriptomic and interaction-based biomarkers described in the text (See **STAR Methods**), calculated for all non-ICB targeted therapies (N = 512). **(d)** OR (y-axis) of ENLIGHT (blue) and the other biomarkers detailed in panel **c**, as well as SELECT and three ICB-specific transcriptomic biomarkers, calculated for the ICB cohorts (N = 161). Biomarkers denoted by ‘X & Y’ (e.g. ‘ENLIGHT & IFN-γ’) were calculated as the geometric mean of X and Y. ‘Predictive biomarker’: a biomarker that is drug-specific, i.e., yields a different prediction per drug. ‘Prognostic biomarker’: a biomarker that is drug-agnostic, i.e., it is predictive of patient survival/general inclination of response but can not be used to guide therapy. ‘Predictive biomarker - ICB specific’: a biomarker that is used to predict general response to immunotherapy but without differentiation between treatments. Asterisks represent the corrected p value for a one-sided test of an OR greater than 1: * p<0.1; ** p<0.05.

We next turned to compare ENLIGHT’s performance to (a) that of its predecessor, SELECT, (b) several known transcriptomics-deduced metrics (IFN-γsignature ^17^, proliferation signature ^64^, cytolytic index ^65^, and the drug target expression levels) and, finally (c) several GI-based scores. This comparison was done comprehensively on all the 21 evaluation cohorts (see **STAR Methods**). Finally, (d) we added a comparison to three ICB specific biomarkers (exhausted T-cell ^66^, TIDE ^16^ and CD8+T-cell abundance ^67^) on the immunotherapy datasets. The performance of ENLIGHT could not be compared to that achieved by DNA-based markers such as TMB or MSI, since all evaluation datasets available include only mRNA data. We note that several of the widely used biomarkers included in these comparisons do not provide differential scoring for different drugs, i.e., they cannot be used to guide therapy (e.g. proliferation signature), while others are specific to predicting response to ICB (e.g. exhausted T-cell). Nevertheless, all are included for completeness.

As all biomarkers other than SELECT and TIDE do not suggest a threshold for determining favorable treatments, we set the decision threshold for those using the tuning sets to match ENLIGHT’s recall (see **STAR Methods**). For TIDE, we used the threshold of 0, as suggested in Jiang et al. ^16^ and for SELECT, we used 0.44 as suggested in Lee et al. ^29^. For 11 of the 21 evaluation cohorts (N=400), the GI networks produced by SELECT were too small to produce robust predictions (having less than 4 GIs). **Figure 3b** compares the odds ratio of ENLIGHT and SELECT on the remaining 10 cohorts (N = 297). While SELECT provides a beneficial discriminatory power on the evaluation cohorts to which it could be applied, especially for ICBs (**Figure 3d**), ENLIGHT can analyze a wider range of treatments than SELECT, with overall higher precision. **Figures 3c and 3d** compare ENLIGHT’s performance to all the biomarkers enumerated above. This is presented separately for non-ICB cohorts (N = 512, the *Selinexor* dataset was excluded since IFN-γ, cytolytic index and target expression could not be calculated on this dataset due to missing data) and for the ICB cohorts (N = 152, the *Anti-PD1 +-Anti-CTLA4* dataset was excluded since missing data prevented calculation of the proliferation signature), since three of the markers are ICB-specific, and since SELECT could only be applied to a few of the non-ICB datasets, as described above. **Table S4** contains the detailed statistics of all comparisons. In short, ENLIGHT has significantly higher OR than SELECT on the datasets for which SELECT could produce predictions (**Figure 3b**, p = 0.0487). In addition, ENLIGHT has higher OR than all other markers, with the improvement being statistically significant in all non-ICB datasets (**Table S4**).

In light of the high predictive power of several ICB-specific markers (namely, proliferation signature, cytolytic index and IFN-γ signature), we studied if ENLIGHT’s combination with these biomarkers can further enhance the overall predictive power for response to ICB. To this end, we tested the predictive power of new combined biomarkers computed as the geometric mean of: (i) ENLIGHT and each of the three individual markers; (ii) ENLIGHT and all three biomarkers; and (iii) the three markers without ENLIGHT (for comparison). For each of these combined biomarkers, we optimized the decision threshold as explained in **STAR Methods.** We found that the combination of ENLIGHT with each of the three biomarkers is more predictive than either ENLIGHT or any of them on their own, with the combination of ENLIGHT and IFN-γbeing superior to all other combinations and individual markers, achieving an OR of 4.07 in the ICB cohorts (**Figures 3d and S4a**). Notably, in the non-ICB cohorts ENLIGHT on its own is superior to the combined biomarkers (**Figures 3c and S5**).

Overall, these results provide strong support for the ability of ENLIGHT to provide clinical benefit in the precision medicine scenario with superior performance over known markers, spanning a variety of different treatments and cancer types.

### ENLIGHT Enables Near Optimal Exclusion of Non-responding Patients in Clinical Trial Design

In the Clinical Trial Design scenario, we are interested in identifying a sub-population of non-responding patients who could be excluded from the trial a priori, thereby allowing smaller studies to achieve higher response rates with adequate statistical power. **Figure 4**(top row) depicts the proportion of true non-responders among those predicted not to respond (NPV) as a function of the percent of patients excluded, where patients are excluded by order of increasing EMS. For both Immunotherapy and other mAbs, ENLIGHT’s NPV curve is considerably higher than the NPV expected by chance, i.e., the percentage of non-responders, testifying to its benefit. For targeted small molecules, however, it is unable to reliably identify non-responders, an issue that should be further studied and improved upon in future work.

**Figure 4:**
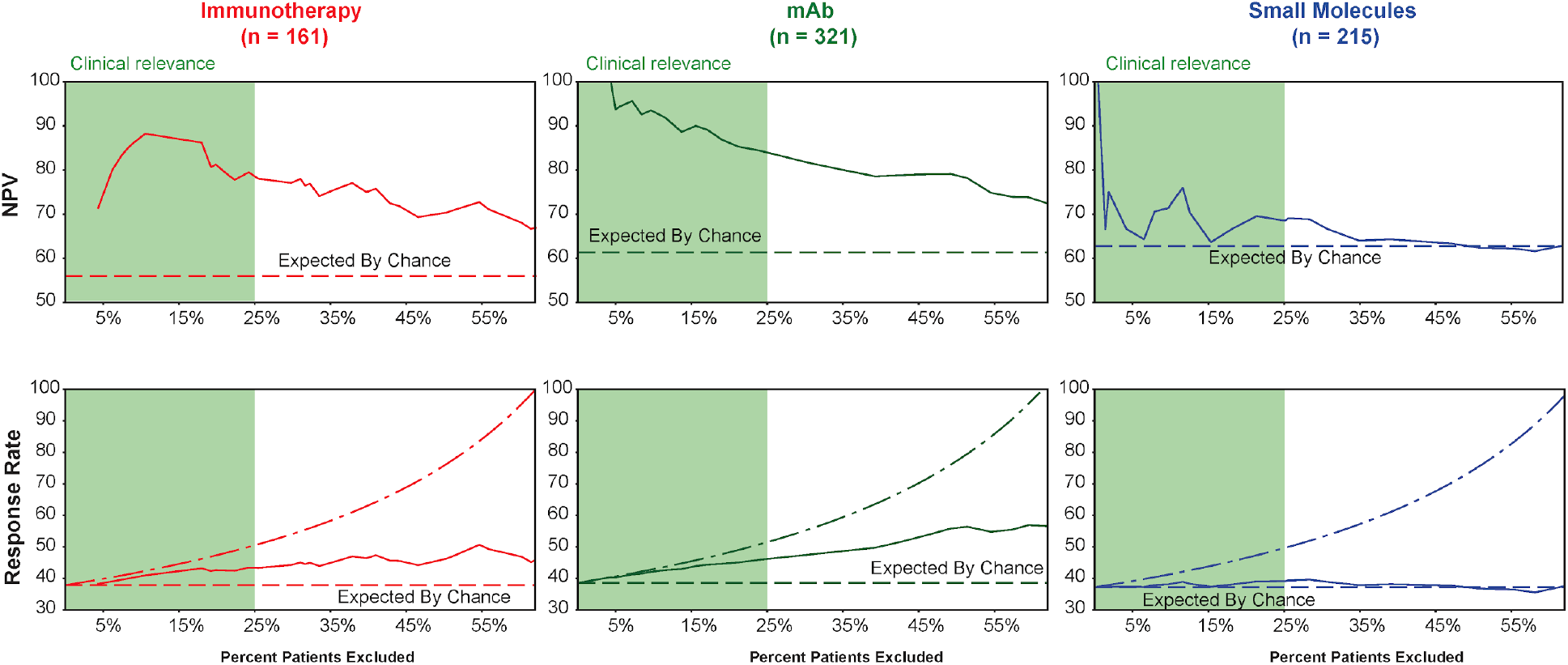
ENLIGHT can facilitate the exclusion of non-responding patients in Clinical Trials. Each of the three columns depicts ENLIGHT’s results on the aggregate of all evaluation cohorts from a given therapeutic class. **Panels on the top row** display the NPV (percent of true non-responders out of those predicted as non-responders) as a function of the percentage of patients excluded. The horizontal line denotes the actual percent of non-responders in the corresponding aggregate cohort (i.e., the NPV expected by chance). **Panels on the bottom row** display the response rate among the remaining patients (y axis) after excluding a certain percentage of the patients (x axis). The horizontal line denotes the overall response rate in the aggregate cohort. The dotted-dashed line represents the upper bound on the response rate, achieved by the “all knowing” optimal classifier excluding only *true non-responders*.

The bottom row of **Figure 4** depicts the response rate in the remaining cohort after excluding patients with EMS below the decision threshold. As evident, ENLIGHT-based exclusion considerably increases the response rate among the remaining patients (middle, solid line). The dotted-dashed line represents the limit performance of an optimal “all-knowing” classifier that excludes all non-responders, retaining only true responders (correspondingly, the x axes end when this optimal classifier excludes all true non-responders, achieving the optimal response rate of 100%). Focusing on a practical exclusion range of up to 25% of the patients (shaded area), ENLIGHT-based exclusion achieves 87%-97% and 90%-99% of the optimal exclusion response rate, for both Immunotherapy and other mAbs, respectively – see **Table 1**. It is important to acknowledge that the ENLIGHT-based exclusion strategy assumes knowledge of the EMS distribution in the trial, which may not be known a priori, but could be estimated using historical transcriptomics data from a reference population of the pertaining cancer indication and clinical characteristics.

**Table 1:**
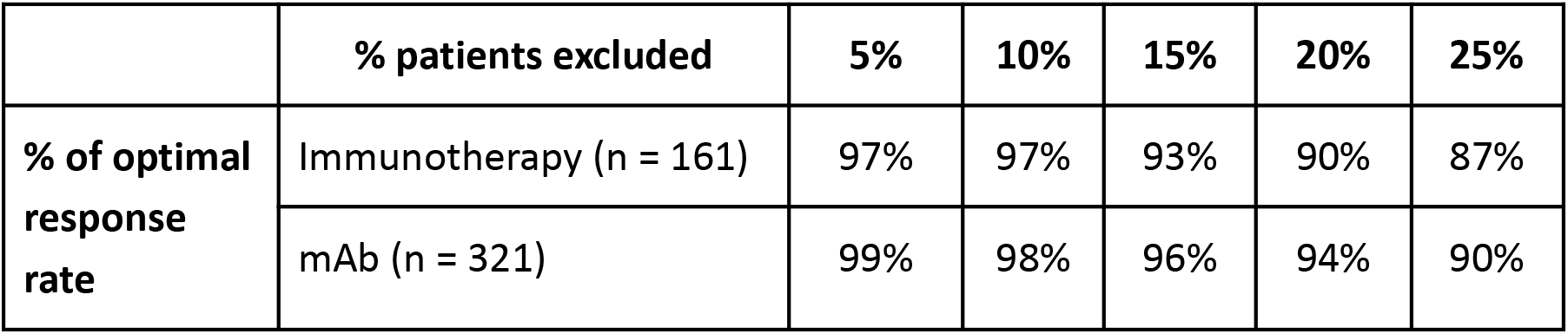
The performance of ENLIGHT’s exclusion strategy in Clinical Trial Design compared to the optimal upper bound. For each percent of patient exclusion (columns), the response rate among the remaining patients when excluding based on increasing EMS, is given as a percentage of the upper bound response rate achieved by the “all knowing” optimal classifier that excludes only *true non-responders*.

|  | % patients excluded | 5% | 10% | 15% | 20% | 25% |
| --- | --- | --- | --- | --- | --- | --- |
| % of optimal response rate | Immunotherapy (n = 161) | 97% | 97% | 93% | 90% | 87% |
|  | mAb (n = 321) | 99% | 98% | 96% | 94% | 90% |

## Discussion

Here we present ENLIGHT, an algorithmic platform that leverages synthetic lethality and rescue interactions to predict response to targeted therapies and immunotherapies. ENLIGHT is an unsupervised approach that, like its predecessor, SELECT, leverages large-scale data in cancer to infer GI networks associated with drug targets on a whole genome scale, and then uses the activation patterns of the genes comprising these networks, as measured in the tumor, to generate a *matching score* for each possible treatment. In difference from its predecessor, ENLIGHT has been designed and evaluated with two real-world clinical scenarios in mind: personalized oncology, where one matches the best treatment to a patient based on a fixed decision threshold, and clinical trial design, where the goal is to a priori exclude non-responding patients in the best possible manner. Testing ENLIGHT on 21 unseen clinical cohorts showed that patients whose treatments were recommended by ENLIGHT have markedly better odds of response than the others. ENLIGHT is a promising systematic approach that, in a comprehensive retrospective analysis, is shown to provide clinically relevant benefits across a broad array of treatments and indications. Despite this broad applicability and its general unsupervised nature, its performance is comparable to that of supervised classifiers trained and predictive on very narrow and specific treatment cohorts. This comparison is of much interest since, in theory, given sufficiently large datasets, such supervised methods are expected to yield higher performance on unseen datasets than unsupervised methods like ENLIGHT. However, if the training sets available are small (as is regrettably the case with the vast majority of the data currently available), practice may differ from theory and supervised methods may overfit and underperform.

ENLIGHT is also more predictive than its predecessor, SELECT, and other transcriptomics-based biomarkers. Some of the biomarkers in these comparisons are prognostic in nature, i.e., provide information about the patient’s overall cancer outcome, regardless of therapy (e.g. proliferation signature) or are specific to predicting response to ICB (e.g. T-cell exhaustion). Unlike ENLIGHT, such markers cannot add information when one aims to choose between various treatments. Specifically, IFN-γhad an odds ratio comparable to that of ENLIGHT on the ICB datasets, which are almost exclusively anti-PD1. However, unlike ENLIGHT, it will not be able to offer differential scoring as more ICB treatments enter the clinic, since its score is not treatment-dependent. The combination of ENLIGHT and IFN-γhas increased predictive performance for ICB therapy, obtaining an impressive OR of 4, and offering differential scoring per treatment, due to ENLIGHT’s predictive nature. Finally, ENLIGHT can enhance clinical trial design, by efficiently excluding non-responsive patients while attaining more than 90% of the maximum attainable response rate in targeted mAbs and ICB cohorts.

In the past several years, bulk RNA-seq has become increasingly available and reliable, opening the door for translational and clinical applications ^68^. The Pan-Cancer Analysis of whole genomes demonstrated how the majority of classical driver genes exhibit alterations that were potentially better characterized via RNA than DNA ^69^. Beyond developing drugs for new targets, we must consider ways to expand eligibility criteria for existing targeted therapies and immunotherapies. Data from ENLIGHT can provide a biologically informed basis to test strongly prioritized off-label therapies for specific patient populations through Phase II trials. This may be especially helpful in rare cancer types or challenging clinical scenarios, where hypothesis generation will only require access to relevant RNA-seq samples. Second, building on the “Clinical Trial Design” capacity of ENLIGHT, the tool can offer simulations focused on a given therapeutic option and its performance across multiple types of patient populations—a computational version of the resource-challenging yet exciting basket trial design.

The current study mainly establishes ENLIGHT’s power to predict monotherapies. However, in clinical practice, identifying combination therapy is often desirable. Notably in this context, ENLIGHT shows promising results when predicting response to targeted therapy on the background of chemotherapy, suggesting that a clinician could use ENLIGHT to identify favorable immune/targeted therapy and combine it with chemotherapy as deemed fit clinically.

While ENLIGHT’s performance is evaluated here on a broad array of unseen clinical trial datasets, its performance should obviously be further evaluated in prospective studies. Indeed, based on the results obtained so far by SELECT and ENLIGHT, the design of such a multi-arm study – SYNTHESIS – is now underway at the National Cancer Institute’s Center for Cancer Research. We hope that the results presented here will elicit additional prospective studies in other clinical centers across the globe and we are committed to facilitating such studies.

### Limitations of study

Like any response prediction approach, ENLIGHT has several limitations that should be acknowledged. First, as it operates on the drug target level, it has very limited utility in predicting response to chemotherapies, and more generally, its prediction accuracy depends on the accuracy with which the targets of a given drug have been identified. Second, most datasets included in this study are based on mRNA expression derived from microarrays, while the standard is shifting towards RNAseq data. Third, current applications of ENLIGHT have focused on bulk tumor transcriptomics. Future work is needed to study its application for analyzing single-cell tumor transcriptomics, to build predictors that consider tumor heterogeneity and the important interplay between a tumor and its immune microenvironment. Fourth, while gene expression is at times strongly associated with protein levels, in other cases it is not, especially for lowly expressed genes. Notably, some of these limitations may point us to future directions for building even better genetic interaction-based predictive approaches. This includes approaches for inferring and harnessing cell-type specific interactions from single cell and deconvolved expression data, approaches that chart the genetic interaction landscape at different regions of the tumor, and more. Finally, one may reasonably anticipate that the advent of much larger molecular and response datasets will on its own contribute to the accuracy of the methods already extant.

## Supporting information

STAR methods

## Author Contributions

G.D., T.B., E.R. and R.A. conceptualized the work described in the study. G.D., E.D.S., Ef.E., E.M., I.M., T.B. and R.A. developed the statistical and algorithmic methodology. G.D., D.S.B.Z. and Em.E. performed data collection and curation. D.J. contributed one of the datasets analyzed (Alpelisib/Ribociclib). G.D., E.D.S., Ef.E., D.S.B.Z., O.T., D.T.H., S.S., N.U.N., J.S.L., T.B., E.R. and R.A. analyzed the data. Z.R., D.J., A.A., W.L.D., S.L., R.B., R.K., A.P.S., F.K., M.R.G., K.A. and P.S.R supported the clinical analyses. B.T. and P.V. developed the software, A.S. tested it. G.D., E.D.S., P.S.R., T.B., E.R. and R.A. wrote the paper. E.S., A.S., Z.R., S.L.M A.P.S., M.R.G. and K.A. made significant edits. All authors read and approved the final article and take responsibility for its content.

## Acknowledgments

This research was supported in part by the Israeli Innovation Authority. This research was supported in part by the Intramural Research Program, National Institutes of Health, National Cancer Institute. The content of this publication does not necessarily reflect the views or policies of the Department of Health and Human Services, nor does mention of trade names, commercial products, or organizations imply endorsement by the U.S. Government.

