## Supplementary material for "Clinically oriented prediction of patient response to targeted and immunotherapies from the tumor transcriptome": STAR methods

### RESOURCE AVAILABILITY

#### Lead contact

#### Materials availability

This study did not generate new unique reagents.

#### Data and code availability

- All transcriptomic and clinical information including treatment and outcome information of the datasets used in the paper are publicly available as individually described in **Table S1**. For ease of usage, all datasets analyzed in this manuscript can be found in https://github.com/PangeaResearch/enlight-data.
- Any additional data required to reanalyze the data reported in this paper is available from the lead contact upon request.
- A web service that generates EMS for the drugs described in this study as well as for any new data supplied by the user is available at https://ems.pangeabiomed.com/

### METHOD DETAILS

#### Data Collection

We surveyed the public domain for available cohorts of patients receiving targeted therapies or immunotherapies, containing both pre-treatment transcriptomics and response information (either RECIST or a binary classification of response). We identified a total of **23 real world datasets** which were not previously analyzed by either SELECT or ENLIGHT, and can hence serve as unseen datasets: 22 datasets (Ascierto et al., 2016; Atwood et al., 2015; Birkbak et al., 2018; Bossi et al., 2016; Byers et al., 2013; Cui et al., 2021; Dieci et al., 2016; Foà et al., 2014; Hsu et al., 2021; Lassman et al., 2022; Liu et al., 2012; Magbanua et al., 2021; Pentheroudakis et al., 2014; Prat et al., 2014; Pusztai et al., 2021; Raponi et al., 2007, 2008; Sammut et al., 2022; Shen et al., 2012; Verstraete et al., 2015; Watanabe et al., 2011; Zhao et al., 2019) from GEO, ArrayExpress, CTRDB or the broader literature published by February 2022, and one dataset that was obtained as part of a collaboration with Massachusetts General Hospital (MGH) which we publish here for the first time. We selected **six** datasets (Dieci et al., 2016; Guarneri et al., 2015; Kakavand et al., 2017; Pinyol et al., 2019; Riaz et al., 2017; Rizos et al., 2014) already analyzed in Lee et al. (Lee et al., 2021) along with **two** (Prat et al., 2014; Watanabe et al., 2011) of these 23 unseen sets to serve as *tuning sets*. These **eight tuning datasets** were selected as they span a range of different treatments, therapeutic classes, response rates and sample sizes, reflecting diverse real-world data, covering five targeted therapies and one immune checkpoint blockade (ICB). These datasets were used to tune the parameters of ENLIGHT, including the GI network size and a decision threshold on the ENLIGHT Matching Score (EMS) that is used for predicting response (see below). We set aside the remaining **21 unseen datasets** – *the evaluation sets* – as unseen data for evaluation.

Table S1 details the tuning and evaluation datasets used in this study. All datasets were coupled with response to treatment in the form of either: (i) RECIST criteria response evaluations or (ii) binary classifications of responders and non-responders that was not exclusively defined using RECIST and in several cases was not specified. In this study, we classify a patient as **responder** if he/she had a RECIST evaluation of CR/PR, or if he/she had a binary classification of responder. The rest were classified as **non-responders**. In each dataset we only analyzed patients for whom both pre-treatment transcriptomics and response data was available. N in the table denotes the number of patients that were analyzed. The *Selinexor* dataset encompasses data from a clinical trial containing several arms, each with a different treatment dosage which significantly affected the response rate. Here we only analyzed the largest arm of the trial (N = 24). In cohort *Anti-PD*$1_4$, which is a combination of immunotherapy and a targeted drug, we analyzed the score based on the immunotherapy only.

**The ENLIGHT Pipeline**

As explained and detailed in the main text and in **Supplementary Note 1**, ENLIGHT improves and extends upon SELECT, making important changes to the pipeline to improve and focus it on translational aspects. For ease of reading, we include here the entire ENLIGHT pipeline, including those parts which are taken from (Lee et al., 2018, 2021; Sahu et al., 2019), and properly refer herein.

The ENLIGHT algorithm can be broadly divided into two steps (see **Figure 1a** for a visualization of the pipeline): (i) the *inference engine* and (ii) the *prediction engine*.

***Inference Engine***

The inference engine uses 4 statistical tests to identify interacting gene pairs:

1. in-vitro test

   By definition, it is expected that gene A will be more essential when its SL partner gene B is inactive in a cancer cell line. Using a set of input genome-wide shRNA/siRNA/sgRNA screens mined from DepMap (Dempster et al.; Meyers et al., 2017; Pacini et al., 2021), ENLIGHT identifies pairs that show **conditional essentiality**: Gene A is defined as *SL conditionally essential* with gene B if its essentiality is significantly higher in cell-lines where gene B is inactive using Wilcoxon rank sum test. We use both mRNA expression and SCNA data to classify genes as over/under-active. Similarly, gene A is defined as *SR-DU/SR-DD conditionally essential* with gene B if its essentiality is significantly higher in the samples where gene B is underactive/overactive. An SR-DU/SR-DD interaction between gene A and gene B dictates that a cell can be **rescued** from cell death caused by the inhibition of gene A by the upregulation/downregulation of gene B respectively.

2. Depletion test

   ENLIGHT follows the ideas of Lee et al. *(Lee et al., 2018)* and Sahu et al. *(Sahu et al., 2019)* in requiring SL/SR pairs not to display joint activation patterns that are disadvantageous for SL/SR interactions in patient tumors. This requirement is implemented by a **depletion test**, added as a step to the inference engine. Conceptually, if an SL interaction exists between a pair of genes, we would expect not to observe tumors in which both genes are inactive, since this would have caused tumor cell lethality. Similarly, we would not expect patients with inactivation of a gene to have low/high activation of its SR-DU/SR-DD rescuer, since that would induce tumor cell death. To summarize, in both SL/SR cases we are looking for the statistical absence (or **depletion**) of a non-favorable joint activation pattern in observational cohorts to support a GI between gene pairs.

3. <u>Survival test</u>

This test identifies candidate SL/SR pairs that confer favorable/unfavorable patient survival when the interaction is active. When an SL pair is simultaneously inactive in a patient or a cell-line we term it **active**. Similarly, we term an SR-DU/SR-DD as active when gene A is inactive in conjunction with its SR-DU/SR-DD partner being underactive/overactive. The survival test used in ENLIGHT is a fully parametric test, based on an exponential survival model. Given the omics of a gene pair from a patient cohort, coupled with survival data, we first calculate a covariate value for each patient, reflecting its joint activation state of the gene pair. For a true SL/SR pair, patients in whom the joint activation of the pair is in a disadvantageous state in the tumor, are expected to have better survival, since this should lead to tumor cell death. Hence, the covariate value is positively associated with survival time in these cases. The statistical model of the test follows the common assumption that covariates have a log-linear effect on survival times, and the coefficient of the SL/SR covariate reflects whether a putative interaction confers a significant effect on patient survival. The model also controls for patient age, gender and stage as confounding factors. If data is missing for any of these three attributes in a patient, we set it to the mean of all other patients (or the majority in the case of gender). Altogether, the likelihood of observing a population with survival times *t* and covariates matrix *X* is given by:

$$L(t, X, \beta) = \prod_{i \in OBS} S\left(t_i, x_i, \beta\right) * \lambda(t_i, x_i, \beta) \prod_{i \notin OBS} S\left(t_i, x_i, \beta\right) = \prod_{i \in OBS} e^{x^T\beta} e^{-e^{x^T\beta}t} \prod_{i \notin OBS} e^{-e^{x^T\beta}t}$$

Where *OBS* are the set of deceased patients, *S* is the survival function, λ is the hazard function, $x_i$ are the covariates for patient i, $t_i$ is the time of death or censoring for patient i and β are the coefficients associated with the covariates. We solve this equation to identify the coefficient β associated with the interaction along with its 95% CI and p-value.

4. Phylogenetic test

   The last test identifies SL/SR pairs with high similarity between their phylogenetic profiles, following observations that interacting genes were found to be conserved across different species (Hartwell et al., 1997; Srivas et al., 2016) and following the analysis of Lee et al. (Lee et al., 2018, 2021). This is done by calculating the Euclidean distance between the genetic similarity profiles of two genes A and B across 86 species, while taking into account the baseline phylogenetic distance between the species (adopting the method of Tabach et al. (Tabach et al., 2013)).

In order to build a GI network around specific drug target/s, we start by performing the above 4 tests for all putative SL/SR-DU/SR-DD pairs between the targets of the drug and all other genes in the genome. Then, we sequentially filter out non-significant pairs for each test, starting from all pairs and all interaction types, so that only pairs that pass all 4 tests are kept. We follow Lee *et al.* for setting the statistical significance thresholds. Finally, we rank the remaining interactions according to the survival test as it best reflects the clinical impact of the interactions, and use the top K interactions to build the GI network. In this study we tested K = 25, 50, 100, 150 and 200 on the *tuning sets* and selected K = 100 as it achieved the best PPV. For immunotherapies, ENLIGHT uses the same GI networks used in SELECT (Lee et al., 2021) (K = 10).

***Prediction Engine***

The prediction engine predicts the response to a given drug or a given combination of drugs with known drug targets based on quantitative RNA data (microarray or RNAseq). The prediction engine works as follows:

GI network inference

First, a GI network surrounding all drug targets is built based on the inference engine.

Normalization

Next, the RNA data of the cohort is rank normalized to values in [0,1] twice: gene-wise and patient-wise. Gene-wise normalized values are used to identify gene activation states of SL/SR partners across comparable samples of the same tissue. Patient-wise normalized values are used to determine whether the drug targets are expressed to a minimal degree in a patient for the drug to have an effect.

Scoring

The ENLIGHT Matching Score (EMS) is defined as the fraction of SL/SR interactions that are in an advantageous predisposition for drug admission. That is, a drug that inhibits gene A is expected to work better in patients for whom an SL/SR-DU B of A is underexpressed, or for whom an SR-DD partner B of A is overexpressed. Thus, the fraction of under/over expressed partners in advantageous states (with respect to the interaction type) is expected to be positively associated with response. A gene is determined to be underexpressed if its normalized expression is equal to or below 1/3 (i.e. is in the bottom tertile across samples in the same dataset), or overexpressed if its normalized expression is equal to or above 2/3 (i.e. is in the top tertile across samples in the same dataset), similar to Lee et al. *(Lee et al., 2021)*. In addition, we zero the EMS of a patient if its mean patient-wise normalized expression of drug targets is below or equal to the 30th percentile across genes. This has been motivated by the notion that an antagonist drug will not be effective when its target genes are underexpressed. Finally, for treatments that are highly target specific, namely ICB and other mAbs, the EMS incorporates the target expression, since the drug is expected to be more effective when the target expression is higher (Dolled-Filhart et al., 2016; Slamon et al., 2001). Specifically, the EMS is a geometric mean of the network-based score and a logistic function of the target expression. Drug target genes were mapped based on DrugBank (Law et al., 2014).

**Biomarkers compared with ENLIGHT**

In order to compare the OR for identifying beneficial treatments between different biomarkers, the focus and objective of this study, one has to set a clinical threshold for binary response classification. For two of the biomarkers compared in this study, such thresholds were given in the original publications. For SELECT, we used the threshold of 0.44 as described in Lee et al. (Lee et al., 2021) and for TIDE we used the threshold of 0 as suggested by Jiang et al (Jiang et al., 2018). For the remaining markers, we used the tuning sets to assign thresholds. For each biomarker, we calculated its scores in the tuning datasets, and searched for a threshold that matches ENLIGHT's recall:

1. For comparing ENLIGHT to other biomarkers on non-ICB datasets (**Figure 3c**), we identified a threshold for each of the presented biomarkers based on all tuning datasets except *Anti-PD1*.
2. For comparing ENLIGHT to other biomarkers on ICB datasets (**Figure 3d**), we identified a threshold for each of the presented biomarkers based only on *Anti-PD1*.

The difference in threshold identification is intended to make an appropriate comparison to ENLIGHT's recall which naturally varies between the two cases. The identified thresholds were then used to calculate OR based on the test datasets for **Figure 3**.

In order to avoid artifacts arising from varying mRNA quantification platform (i.e. RNAseq vs. microarrays), for example the difference in the dynamic range of expression between platforms, all biomarkers that do not have explicit means for calculation (i.e. all markers except SELECT, TIDE and CD8+ abundance which have available code for calculation) were calculated based on gene-wise ranking of expression across samples, after applying proper normalizations to account for library size and gene lengths (if needed). The description of each marker is as follows:

<u>Target Expression</u>

Mean expression of all drug targets in the treatment regimen.

<u>ENLIGHT-InVitro</u>

An EMS-like score that uses as the GI network, all interactions that passed the first test of ENLIGHT: the In-Vitro test.

Other drugs SL

An EMS-like score that uses the combined GI network of **all** drugs shown in **Figure 2a** except the given drug.

Random Genes

An EMS-like score based on a GI network populated by a set of random genetic interactions. That is, we randomly choose 100 gene partners and an interaction type for each (SL/SR), and use them to score each sample in the same way the EMS is calculated.

IFN-γ

A score based on a signature that measures the expression of the Interferon-gamma pathway (Ayers et al., 2017).

Proliferation

A score based on the proliferation signature suggested by (Whitfield et al., 2006).

Cytolytic

A score based on the Cytolytic index suggested by (Rooney et al., 2015).

Exhausted T-cell

A signature that measure T-cell exhaustion in the TME as suggested by (Wherry et al., 2007).

TIDE

a gene expression-based classifier that predicts response to ICB based on T-cell exclusion/exhaustion developed by (Jiang et al., 2018).

CD8+ T-cell

CD8+ T-cell abundance as calculated by CYBERSORT (Newman et al., 2015).

**QUANTIFICATION AND STATISTICAL ANALYSIS**

Differences between distributions were analyzed using a one-sided Mann-Whitney test using the scipy.stats.mannwhitneyu function in python. Test of OR > 1 was done using Fisher's exact test. Difference between OR was done using a one-sided test for difference in OR. Difference between PPV and overall response rates was done using a one-sided proportion test. Difference between OR of two methods was done using Chi square test. All statistical results were FDR corrected for multiple hypotheses throughout the paper. Throughout the paper, we denoted p <

0.1 with a single asterix (*) and $p < 0.05$ with two asterix (**). Throughout the paper, brackets denote 95% CI and N denotes the sample size of the dataset.

Dempster, J.M., Rossen, J., Kazachkova, M., Pan, J., Kugener, G., Root, D.E., and Tsherniak, A. Extracting Biological Insights from the Project Achilles Genome-Scale CRISPR Screens in Cancer Cell Lines. https://doi.org/10.1101/720243.

Dieci, M.V., Prat, A., Tagliafico, E., Paré, L., Ficarra, G., Bisagni, G., Piacentini, F., Generali, D.G.,

Conte, P., and Guarneri, V. (2016). Integrated evaluation of PAM50 subtypes and immune modulation of pCR in HER2-positive breast cancer patients treated with chemotherapy and HER2-targeted agents in the CherLOB trial. Ann. Oncol. *27*, 1867–1873. .

Dolled-Filhart, M., Roach, C., Toland, G., Stanforth, D., Jansson, M., Lubiniecki, G.M., Ponto, G., and Emancipator, K. (2016). Development of a Companion Diagnostic for Pembrolizumab in Non-Small Cell Lung Cancer Using Immunohistochemistry for Programmed Death Ligand-1. Arch. Pathol. Lab. Med. *140*, 1243–1249. .

Foà, R., Del Giudice, I., Cuneo, A., Del Poeta, G., Ciolli, S., Di Raimondo, F., Lauria, F., Cencini, E., Rigolin, G.M., Cortelezzi, A., et al. (2014). Chlorambucil plus rituximab with or without maintenance rituximab as first-line treatment for elderly chronic lymphocytic leukemia patients. American Journal of Hematology *89*, 480–486. https://doi.org/10.1002/ajh.23668.

Guarneri, V., Dieci, M.V., Frassoldati, A., Maiorana, A., Ficarra, G., Bettelli, S., Tagliafico, E., Bicciato, S., Generali, D.G., Cagossi, K., et al. (2015). Prospective Biomarker Analysis of the Randomized CHER-LOB Study Evaluating the Dual Anti-HER2 Treatment With Trastuzumab and Lapatinib Plus Chemotherapy as Neoadjuvant Therapy for HER2-Positive Breast Cancer. Oncologist *20*, 1001–1010. .

Hartwell, L.H., Szankasi, P., Roberts, C.J., Murray, A.W., and Friend, S.H. (1997). Integrating genetic approaches into the discovery of anticancer drugs. Science *278*, 1064–1068. .

Hsu, C.-L., Ou, D.-L., Bai, L.-Y., Chen, C.-W., Lin, L., Huang, S.-F., Cheng, A.-L., Jeng, Y.-M., and Hsu, C. (2021). Exploring Markers of Exhausted CD8 T Cells to Predict Response to Immune Checkpoint Inhibitor Therapy for Hepatocellular Carcinoma. Liver Cancer *10*, 346–359. .

Jiang, P., Gu, S., Pan, D., Fu, J., Sahu, A., Hu, X., Li, Z., Traugh, N., Bu, X., Li, B., et al. (2018). Signatures of T cell dysfunction and exclusion predict cancer immunotherapy response. Nat. Med. *24*, 1550–1558. .

Kakavand, H., Rawson, R.V., Pupo, G.M., Yang, J.Y.H., Menzies, A.M., Carlino, M.S., Kefford, R.F., Howle, J.R., Saw, R.P.M., Thompson, J.F., et al. (2017). PD-L1 Expression and Immune Escape in Melanoma Resistance to MAPK Inhibitors. Clin. Cancer Res. *23*, 6054–6061. .

Lassman, A.B., Wen, P.Y., van den Bent, M.J., Plotkin, S.R., Walenkamp, A.M.E., Green, A.L., Li, K., Walker, C.J., Chang, H., Tamir, S., et al. (2022). A Phase II Study of the Efficacy and Safety of Oral Selinexor in Recurrent Glioblastoma. Clin. Cancer Res. *28*, 452–460. .

Lee, J.S., Das, A., Jerby-Arnon, L., Arafeh, R., Auslander, N., Davidson, M., McGarry, L., James, D., Amzallag, A., Park, S.G., et al. (2018). Harnessing synthetic lethality to predict the response to cancer treatment. Nat. Commun. *9*, 2546. .

Lee, J.S., Nair, N.U., Dinstag, G., Chapman, L., Chung, Y., Wang, K., Sinha, S., Cha, H., Kim, D., Schperberg, A.V., et al. (2021). Synthetic lethality-mediated precision oncology via the tumor transcriptome. Cell *184*, 2487–2502.e13. .

Liu, J.C., Voisin, V., Bader, G.D., Deng, T., Pusztai, L., Symmans, W.F., Esteva, F.J., Egan, S.E., and Zacksenhaus, E. (2012). Seventeen-gene signature from enriched Her2/Neu mammary tumor-initiating cells predicts clinical outcome for human HER2+:ERα- breast cancer. Proc. Natl. Acad. Sci. U. S. A. *109*, 5832–5837. .

Magbanua, M.J.M., Li, W., Wolf, D.M., Yau, C., Hirst, G.L., Swigart, L.B., Newitt, D.C., Gibbs, J., Delson, A.L., Kalashnikova, E., et al. (2021). Circulating tumor DNA and magnetic resonance imaging to predict neoadjuvant chemotherapy response and recurrence risk. NPJ Breast Cancer *7*, 32. .

Meyers, R.M., Bryan, J.G., McFarland, J.M., Weir, B.A., Sizemore, A.E., Xu, H., Dharia, N.V., Montgomery, P.G., Cowley, G.S., Pantel, S., et al. (2017). Computational correction of copy number effect improves specificity of CRISPR-Cas9 essentiality screens in cancer cells. Nat. Genet. *49*, 1779–1784. .

Newman, A.M., Liu, C.L., Green, M.R., Gentles, A.J., Feng, W., Xu, Y., Hoang, C.D., Diehn, M., and Alizadeh, A.A. (2015). Robust enumeration of cell subsets from tissue expression profiles. Nat. Methods *12*, 453–457. .

Pacini, C., Dempster, J.M., Boyle, I., Gonçalves, E., Najgebauer, H., Karakoc, E., van der Meer, D., Barthorpe, A., Lightfoot, H., Jaaks, P., et al. (2021). Integrated cross-study datasets of genetic dependencies in cancer. Nat. Commun. *12*, 1661. .

Pentheroudakis, G., Kotoula, V., Fountzilas, E., Kouvatseas, G., Basdanis, G., Xanthakis, I., Makatsoris, T., Charalambous, E., Papamichael, D., Samantas, E., et al. (2014). A study of gene expression markers for predictive significance for bevacizumab benefit in patients with metastatic colon cancer: a translational research study of the Hellenic Cooperative Oncology Group (HeCOG). BMC Cancer *14*, 111. .

Pinyol, R., Montal, R., Bassaganyas, L., Sia, D., Takayama, T., Chau, G.-Y., Mazzaferro, V., Roayaie, S., Lee, H.C., Kokudo, N., et al. (2019). Molecular predictors of prevention of recurrence in HCC with sorafenib as adjuvant treatment and prognostic factors in the phase 3 STORM trial. Gut *68*, 1065–1075. .

Prat, A., Bianchini, G., Thomas, M., Belousov, A., Cheang, M.C.U., Koehler, A., Gómez, P., Semiglazov, V., Eiermann, W., Tjulandin, S., et al. (2014). Research-based PAM50 subtype predictor identifies higher responses and improved survival outcomes in HER2-positive breast cancer in the NOAH study. Clin. Cancer Res. *20*, 511–521. .

Pusztai, L., Yau, C., Wolf, D.M., Han, H.S., Du, L., Wallace, A.M., String-Reasor, E., Boughey, J.C., Chien, A.J., Elias, A.D., et al. (2021). Durvalumab with olaparib and paclitaxel for high-risk HER2-negative stage II/III breast cancer: Results from the adaptively randomized I-SPY2 trial. Cancer Cell *39*, 989–998.e5. .

Raponi, M., Harousseau, J.-L., Lancet, J.E., Löwenberg, B., Stone, R., Zhang, Y., Rackoff, W., Wang,

Y., and Atkins, D. (2007). Identification of molecular predictors of response in a study of tipifarnib treatment in relapsed and refractory acute myelogenous leukemia. Clin. Cancer Res. *13*, 2254–2260. .

Raponi, M., Lancet, J.E., Fan, H., Dossey, L., Lee, G., Gojo, I., Feldman, E.J., Gotlib, J., Morris, L.E., Greenberg, P.L., et al. (2008). A 2-gene classifier for predicting response to the farnesyltransferase inhibitor tipifarnib in acute myeloid leukemia. Blood *111*, 2589–2596. .

Riaz, N., Havel, J.J., Makarov, V., Desrichard, A., Urba, W.J., Sims, J.S., Hodi, F.S., Martín-Algarra, S., Mandal, R., Sharfman, W.H., et al. (2017). Tumor and Microenvironment Evolution during Immunotherapy with Nivolumab. Cell *171*, 934–949.e16. .

Rizos, H., Menzies, A.M., Pupo, G.M., Carlino, M.S., Fung, C., Hyman, J., Haydu, L.E., Mijatov, B., Becker, T.M., Boyd, S.C., et al. (2014). BRAF inhibitor resistance mechanisms in metastatic melanoma: spectrum and clinical impact. Clin. Cancer Res. *20*, 1965–1977. .

Rodon, J., Soria, J.-C., Berger, R., Miller, W.H., Rubin, E., Kugel, A., Tsimberidou, A., Saintigny, P., Ackerstein, A., Braña, I., et al. (2019). Genomic and transcriptomic profiling expands precision cancer medicine: the WINTHER trial. Nat. Med. *25*, 751–758. .

Rooney, M.S., Shukla, S.A., Wu, C.J., Getz, G., and Hacohen, N. (2015). Molecular and genetic properties of tumors associated with local immune cytolytic activity. Cell *160*, 48–61. .

Sahu, A.D., S Lee, J., Wang, Z., Zhang, G., Iglesias-Bartolome, R., Tian, T., Wei, Z., Miao, B., Nair, N.U., Ponomarova, O., et al. (2019). Genome-wide prediction of synthetic rescue mediators of resistance to targeted and immunotherapy. Mol. Syst. Biol. *15*, e8323. .

Sammut, S.-J., Crispin-Ortuzar, M., Chin, S.-F., Provenzano, E., Bardwell, H.A., Ma, W., Cope, W., Dariush, A., Dawson, S.-J., Abraham, J.E., et al. (2022). Multi-omic machine learning predictor of breast cancer therapy response. Nature *601*, 623–629. .

Shen, K., Qi, Y., Song, N., Tian, C., Rice, S.D., Gabrin, M.J., Brower, S.L., Symmans, W.F., O’Shaughnessy, J.A., Holmes, F.A., et al. (2012). Cell line derived multi-gene predictor of pathologic response to neoadjuvant chemotherapy in breast cancer: a validation study on US Oncology 02-103 clinical trial. BMC Med. Genomics *5*, 51. .

Slamon, D.J., Leyland-Jones, B., Shak, S., Fuchs, H., Paton, V., Bajamonde, A., Fleming, T., Eiermann, W., Wolter, J., Pegram, M., et al. (2001). Use of Chemotherapy plus a Monoclonal Antibody against HER2 for Metastatic Breast Cancer That Overexpresses HER2. New England Journal of Medicine *344*, 783–792. https://doi.org/10.1056/nejm200103153441101.

Srivas, R., Shen, J.P., Yang, C.C., Sun, S.M., Li, J., Gross, A.M., Jensen, J., Licon, K., Bojorquez-Gomez, A., Klepper, K., et al. (2016). A Network of Conserved Synthetic Lethal Interactions for Exploration of Precision Cancer Therapy. Mol. Cell *63*, 514–525. .

Tabach, Y., Golan, T., Hernández-Hernández, A., Messer, A.R., Fukuda, T., Kouznetsova, A., Liu,

J.-G., Lilienthal, I., Levy, C., and Ruvkun, G. (2013). Human disease locus discovery and mapping to molecular pathways through phylogenetic profiling. Mol. Syst. Biol. *9*, 692. .

Verstraete, M., Debucquoy, A., Dekervel, J., van Pelt, J., Verslype, C., Devos, E., Chiritescu, G., Dumon, K., D'Hoore, A., Gevaert, O., et al. (2015). Combining bevacizumab and chemoradiation in rectal cancer. Translational results of the AXEBeam trial. Br. J. Cancer *112*, 1314–1325. .

Watanabe, T., Kobunai, T., Yamamoto, Y., Matsuda, K., Ishihara, S., Nozawa, K., Iinuma, H., Konishi, T., Horie, H., Ikeuchi, H., et al. (2011). Gene expression signature and response to the use of leucovorin, fluorouracil and oxaliplatin in colorectal cancer patients. Clin. Transl. Oncol. *13*, 419–425. .

Wherry, E.J., Ha, S.-J., Kaech, S.M., Haining, W.N., Sarkar, S., Kalia, V., Subramaniam, S., Blattman, J.N., Barber, D.L., and Ahmed, R. (2007). Molecular signature of CD8+ T cell exhaustion during chronic viral infection. Immunity *27*, 670–684. .

Whitfield, M.L., George, L.K., Grant, G.D., and Perou, C.M. (2006). Common markers of proliferation. Nat. Rev. Cancer *6*, 99–106. .

Zhao, J., Chen, A.X., Gartrell, R.D., Silverman, A.M., Aparicio, L., Chu, T., Bordbar, D., Shan, D., Samanamud, J., Mahajan, A., et al. (2019). Immune and genomic correlates of response to anti-PD-1 immunotherapy in glioblastoma. Nat. Med. *25*, 462–469. .
