## Supplementary Material for "Clinically oriented prediction of patient response to targeted and immunotherapies from the tumor transcriptome"

#### Supplementary Notes

##### Note 1. ENLIGHT improves upon SELECT

In developing ENLIGHT, we have extended and improved SELECT, by introducing the following adaptations:

1. ENLIGHT leverages a larger amount of *in-vitro* data that was updated since SELECT was published. This results in a more robust list of initial GI candidate pairs.
2. While SELECT uses 25 SL pairs in its targeted drug GI networks, ENLIGHT's GI networks for targeted therapies include a combined list of 100 SL and/or SR interactions that are concomitantly inferred for each drug. The fact that ENLIGHT utilizes both SL and SR interactions considerably increases the number of drugs for which it can infer a GI network, and hence produce predictions. In addition, using larger networks reduces the variance in score distributions across treatments and cancer types, allowing a uniform test for multiple drugs. For immunotherapies, SELECT had full coverage and hence ENLIGHT uses the same size 10 GI networks.
3. ENLIGHT follows Lee et al. [1] and Sahu et al. [2] in requiring SL/SR pairs to display a low joint disadvantageous/advantageous activation state in clinical samples, which is reflected by a **depletion test**, added as a step in the inference engine (not present in SELECT). In developing ENLIGHT, the depletion test went under complete revision. Instead of a hypergeometric test on categorized data to identify depletion for both SL and SR interactions, ENLIGHT's depletion test differs between SL and SR: for the SL case, the depletion test is built on the fundamentals of the Gumbel copula, applied on continuous RNA expression data to identify pairs with low probability of being simultaneously inactive. The depletion test for SR requires that the activation state of a rescuer gene be conditioned on its partner being inactive.
4. SELECT uses cox proportional hazard test on categorized expression data to select candidate SL/SR pairs that confer favorable/unfavorable patient survival when the

interaction is active. To increase robustness and statistical power, ENLIGHT applies a fully parametric test, based on an exponential survival model, on continuous expression data.

5. For treatments that are highly target specific, namely ICB and other mAbs, the ENLIGHT matching score incorporates the target expression, since the drug is expected to be more effective when the target expression is higher. Specifically, the EMS is a geometric mean of the network-based score and a logistic function of the target expression.

**Note 2. Analysis of the WINTHER trial**

We analyzed 100 patients for whom both treatment outcome and transcriptomic data were available. Of these, 97 received at least one targeted or immunotherapy agent, 1 of which had missing values that deemed the case non-analyzable. Thus, in total, we calculated an ENLIGHT Matching Score for 96 patients . Among the remaining 96 patients, one patient had a complete response and 11 patients had a partial response. These 12 patients were considered responders, while the 15 patients with stable disease and the 69 patients with progressive disease, were considered non-responders. To infer a GI network for a regimen involving several drugs, we considered the union of the targets from all the drugs in the regimen. We did not consider the targets of chemotherapies or hormonal therapies that were part of the treatment.

Figure S3b shows the EMS of each patient for the prescribed regimen (the *Winther* row) as well as for all ENLIGHT-analyzable drugs in the WINTHER trial. We observe that ENLIGHT identified at least one favorable treatment for all but one patient, and thus, in theory, other drugs or drug combinations may have proven more beneficial for those patients who did not respond.

### Supplementary Figures and Tables

| **Tuning cohorts** | | | | | | |
|---|---|---|---|---|---|---|
| **Name** | **Source** | **Indication** | **Analyzable Drug** | **Background Therapy** | **N** | **N responders** |
| *Bevacizumab* | GSE19860 [3] | Colorectal | Bevacizumab | FOLFOX | 12 | 5 |

| | | | | | | |
|---|---|---|---|---|---|---|
| *Sorafenib* | GSE109211 [4] | HCC | Sorafenib | None | 67 | 21 |
| *Lapatinib* | GSE66399 [5] | Breast | Lapatinib | Chemotherapy | 65 | 21 |
| *Trastuzumab* | GSE50948 [6] | Breast | Trastuzumab | AT followed by CMF | 63 | 31 |
| *BRAFi* | GSE65185 [7] | Melanoma | Vemurafenib/Dabrafenib | None | 17 | 14 |
| | GSE99898 [8] | | | | 16 | 10 |
| | GSE50509 [9] | | | | 20 | 14 |
| *Anti-PD1* | GSE91061 [10] | Melanoma | Nivolumab | None | 50 | 10 |
| **Evaluation cohorts** | | | | | | |
| **Name** | **Source** | **Indication** | **Analyzable Drug** | **Background Therapy** | **N** | **N responders** |
| *Bevacizumab*$_2$ | GSE53127 [11] | Colorectal | Bevacizumab | None | 18 | 3 |
| *Bevacizumab*$_3$ | GSE103668 [12] | Breast | Bevacizumab | Platinum | 21 | 7 |
| *Bevacizumab*$_4$ | GSE60331 [13] | Colorectal | Bevacizumab | Chemoradiation | 17 | 8 |
| *Sorafenib*$_2$ | GSE33072 [14] | Breast | Sorafenib | None | 39 | 20 |
| *Trastuzumab*$_2$ | GSE66399 [5] | Breast | Trastuzumab | Chemotherapy | 23 | 6 |
| *Trastuzumab*$_3$ | GSE37946 [15] | Breast | Trastuzumab | Chemotherapy | 50 | 27 |
| *Trastuzumab*$_4$ | GSE42822 [16] | Breast | Trastuzumab | FEX/TX | 25 | 12 |
| *Trastuzumab*$_5$ | Sammut et al .[17] | Breast | Trastuzumab | FEC | 65 | 19 |
| *Cetuximab* | GSE65021 [18] | H&N | Cetuximab | Platinum | 40 | 14 |
| *Selinexor* | GSE186332 [19] | GBM | Selinexor | None | 24 | 8 |
| *MK*2206 | GSE150576 [20] | Breast | MK2206 | None | 20 | 6 |
| *Tipifarnib*$_1$ | GSE5122 [21] | AML | Tipifarnib | None | 57 | 13 |
| *Tipifarnib*$_2$ | GSE8970 [22] | AML | Tipifarnib | None | 34 | 13 |
| *Rituximab* | GSE35935[23] | CLL | Rituximab | Chlorambucil | 62 | 16 |
| *Alpelisib/Ribociclib* | This manuscript | Breast | Alpelisib/Ribociclib | None | 28 | 16 |
| *Anti-PD1*$_2$ | Zhao et al. [24] | GBM | Nivolumab/Pembrolizumab | None | 15 | 9 |
| *Anti-PD1 +- Anti-CTLA4* | GSE140901 [25] | HCC | Nivolumab/Nivolumab + Ipilimumab/PDR001 + MBG45 | None | 9 | 5 |
| *Anti-PD1*$_3$ | GSE67501 [26] | RCC | Nivolumab | None | 11 | 4 |
| *Anti-PD1*$_4$ | GSE173839 [27] | Breast | Durvalumab | Olaparib | 71 | 29 |
| *Anti-PD1*$_5$ | Cui et al. [28] | Melanoma | Anti-PD1 (drug undisclosed) | None | 55 | 14 |

| | | | | | | |
|---|---|---|---|---|---|---|
| *Vismodegib* | GSE58375 [29] | BCC | Vismodegib | None | 13 | 4 |

**Table S1**. Cohorts used for tuning (top) and evaluation (bottom). Name: Name as appears in main text. Source: The source from which the datasets were obtained. All datasets are also available in https://github.com/PangeaResearch/enlight-data. Analyzable Drug: the drug used for ENLIGHT score generation. 'A/B' indicates a patient received either A or B. 'A +- B' indicates that some patients received A and some received both A and B. Background Therapy: treatments used in combination with the analyzable drug that were not considered when calculating the EMS. N: number of patients analyzed. N responders: number of patients classified as responders.

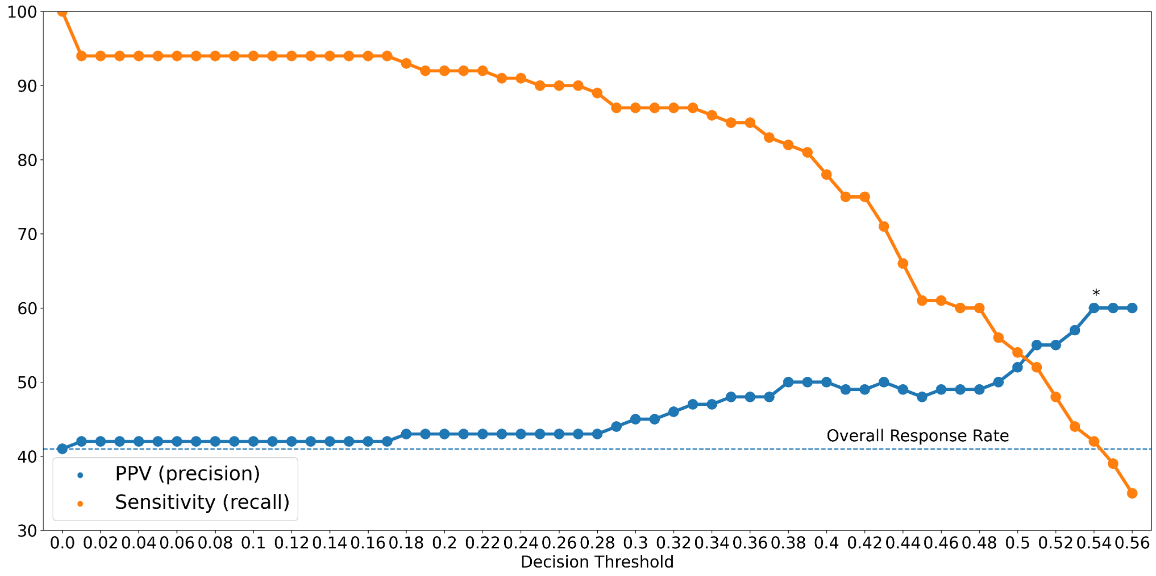


**Figure S1. PPV (precision) and sensitivity (recall) for EMS on the tuning cohorts**. The y axis displays the PPV or sensitivity as a function of the decision threshold shown on the x axis. Figure depicts decision thresholds for which there was at least 30% recall.

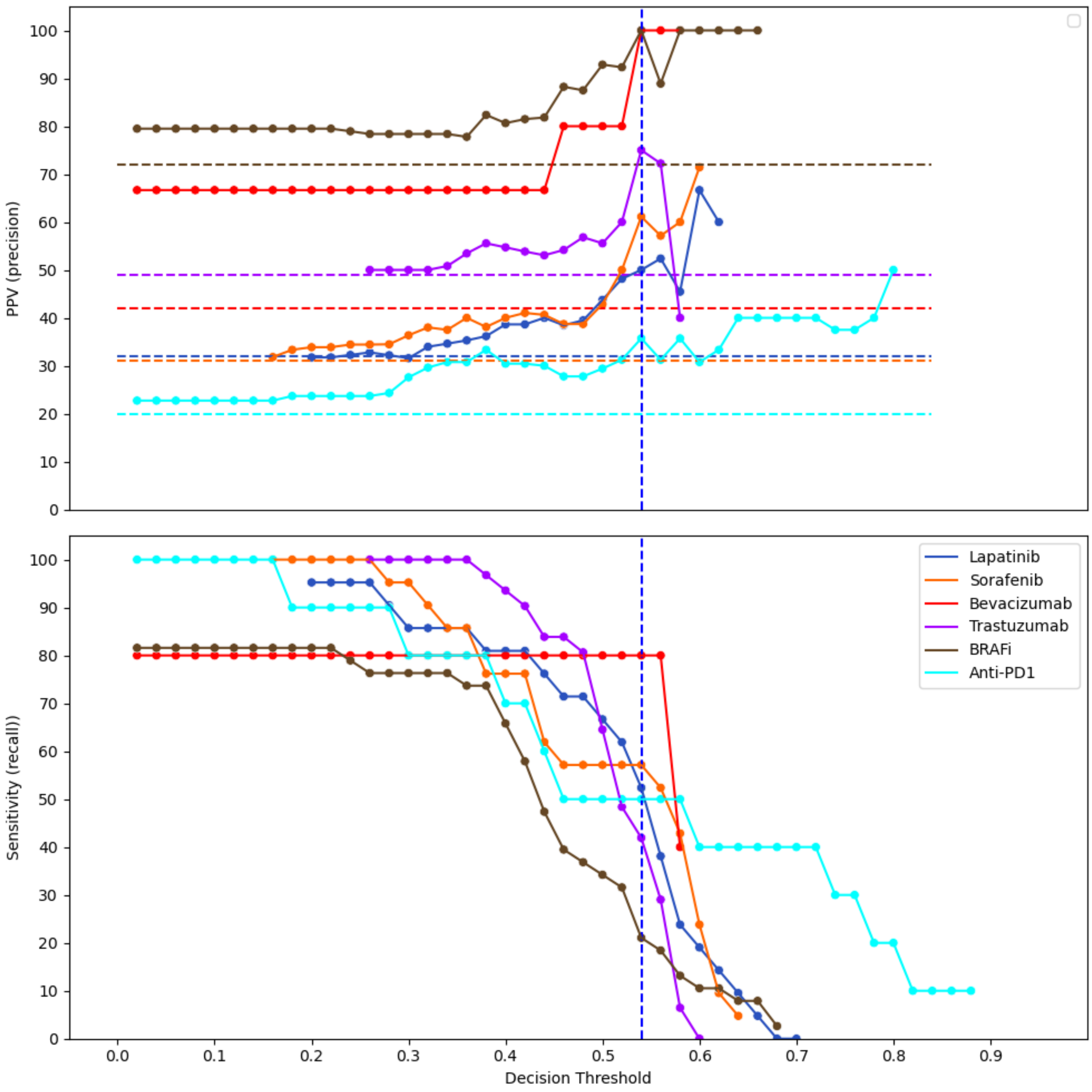


**Figure S2. PPV and Recall as a function of the decision threshold**. The PPV (precision) (top) and sensitivity (recall) (bottom) are presented separately for each of the tuning cohorts. Horizontal lines of corresponding color depict the baseline PPV, i.e., the overall response rate in the cohort.

| | N | Response Rate | PPV (precision) | OR |
|---|---|---|---|---|
| Targeted Small Molecules | 215 | 37% | 45%, *p*=0.018 | 1.59 [0.74, 2.45], *p*=0.094 |
| ICB | 161 | 38% | 49%, *p*=0.0008 | 2.39 [0.98, 4.84], *p*=0.007 |
| mAb | 321 | 39% | 57%, *p*=2.37*e*-10 | 3.72 [2.31, 5.98], *p*=2.24*e*-7 |
| All | 697 | 38% | 52%, *p*=3.30*e*-13 | 2.59 [1.85, 3.6], *p*=3.41*e*-8 |

**Table S2**. ENLIGHT Performance by Therapeutic Class. PPV (precision): The percent of patients identified as matching a treatment that indeed responded. The p-values for the difference between PPV and response rate were calculated using the one sample proportion test. Odds Ratio: The odds ratio for response of ENLIGHT-matched cases; Square brackets indicate the 95% confidence interval; The p-value is calculated using Fisher's exact test.

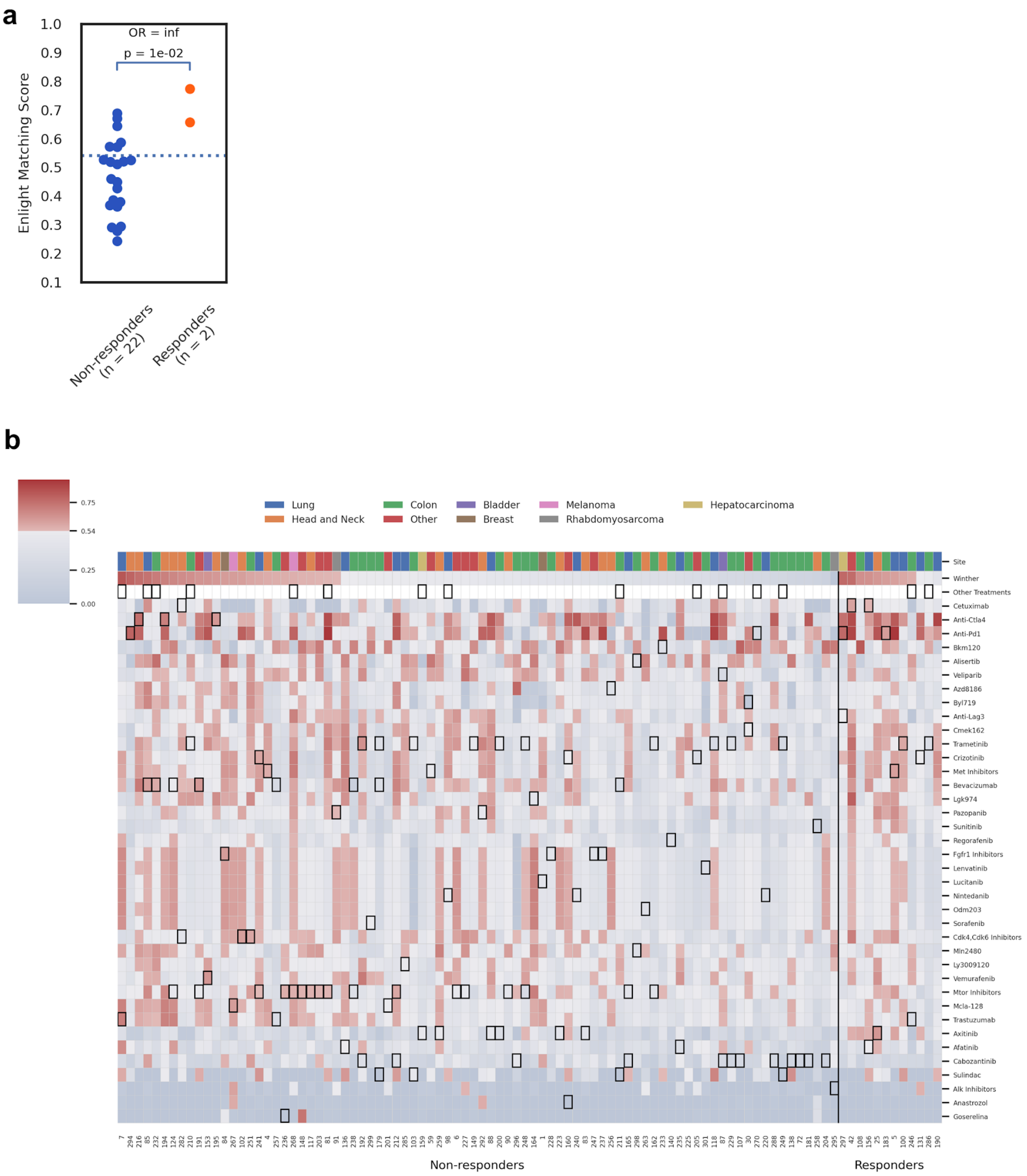
a
OR = inf
p = 1e-02
Enlight Matching Score
1.0
0.9
0.8
0.7
0.6
0.5
0.4
0.3
0.2
0.1
Non-responders (n = 22)
Responders (n = 2)
b
0.75
0.54
0.25
0.00
Lung
Head and Neck
Colon
Other
Bladder
Breast
Melanoma
Rhabdomyosarcoma
Hepatocarcinoma
Site
Winther
Other Treatments
Cetuximab
Anti-Ctla4
Anti-Pd1
Bkm120
Alisertib
Veliparib
Azd8186
Byl719
Anti-Lag3
Cmek162
Trametinib
Crizotinib
Met Inhibitors
Bevacizumab
Lgk974
Pazopanib
Sunitinib
Regorafenib
Fgfr1 Inhibitors
Lenvatinib
Lucitanib
Nintedanib
Odm203
Sorafenib
Cdk4,Cdk6 Inhibitors
Mln2480
Ly3009120
Vemurafenib
Mtor Inhibitors
Mcla-128
Trastuzumab
Axitinib
Afatinib
Cabozantinib
Sulindac
Alk Inhibitors
Anastrozol
Goserelina
Non-responders
Responders

**Figure S3**. (**a**) Analysis of the 24 patients that were treated with a combination of ENLIGHT-analyzable drugs in the WINTHER trial. Responders (orange) have significantly higher EMS than non-responders (blue), p-value is based on one sided Mann-Whitney test. The horizontal line marks the decision threshold for considering a treatment as favorable for a patient ($EMS \geq 0.54$). OR: odds ratio for response for patients receiving treatments with an EMS above the decision threshold. (**b**) The heatmap shows the EMS for the 96 patients analyzed in the WINTHER trial (columns) and all ENLIGHT analyzable drugs given in the trial (rows). The 'Winther' row shows the EMS for the treatment regimen given in the trial. Color designates EMS, with red colors corresponding to ENLIGHET-matched treatments (EMS ≥ 0.54). Black boxes indicate the drugs that were given to each patient. 'Other treatments': non-analyzable drugs, i.e., chemotherapy or hormonal therapy. The cancer type of each sample is color-coded at the top of the heatmap.

| | N | OR |
|---|---|---|
| All | 96 | 11.15 [2.28, 54.54], *p*=7.80e-04 |
| ENLIGHT Analyzable Only | 81 | 10.20 [1.99, 52.24], *p*=2.48e-03 |
| ENLIGHT Analyzable + Unanalyzable | 15 | inf |
| Monotherapy ENLIGHT Analyzable Only | 60 | 8.14 [1.47, 45.18], *p*=0.01 |
| Monotherapy ENLIGHT Analyzable + Unanalyzable | 12 | inf |
| Combination ENLIGHT Analyzable Only | 21 | inf |
| Combination ENLIGHT Analyzable + Unanalyzable | 3 | NA |
| Monotherapy | 72 | 8.40 [1.63, 43.24], *p*=6.07e-03 |
| Combination | 24 | inf |

**Table S3**. ENLIGHT predictions on combination therapies in the WINTHER trial. Patients in the WINTHER trial received either a single drug (monotherapies) or a combination of drugs. Moreover, some of the patients received drugs unanalyzable by ENLIGHT, i.e., chemotherapies or hormonal therapies. N denotes the number of patients; in square brackets - 95% confidence interval for the odds ratio; the p-value is calculated using Fisher's exact test.

| SELECT sets, N = 297 (Figure 3b) | | | | |
|---|---|---|---|---|
| Biomarker | OR | p | 95% CI | p value under H0: ENLIGHT does not have a greater OR than the respective marker (one sided) |
| ENLIGHT | 2.308 | 0.0004 | [1.42,3.73] | |
| SELECT | 1.168 | 0.154 | [0.72,1.875] | 0.0487 |
| non-ICB targeted therapies, N = 512 (Figure 3c) | | | | |
| Biomarker | OR | p | 95% CI | p value under H0: ENLIGHT does not have a greater OR than the respective marker (one sided) |
| ENLIGHT | 2.71 | 1.32E-06 | [1.24,4.59] | |
| gm(ENLIGHT,IFNG) | 2.163 | 0.002 | [1.4,3.341] | 0.233 |
| Proliferation | 0.911 | 1 | [0.636,1.304] | 1.22E-05 |
| IFNG | 1.392 | 0.064 | [0.962,2.014] | 0.006 |
| Cytolytic | 1.44 | 0.064 | [1.003,2.002] | 0.007 |
| gm(IFNG,Cytolytic,Proliferation) | 1.33 | 0.073 | [0.927,1.912] | 0.003 |
| Target Expression | 1.399 | 0.064 | [0.977,2.002] | 0.005 |
| Random Genes | 0.773 | 1 | [0.531,1.125] | 1.72E-06 |
| ENLIGHT-InVitro | 0.824 | 1 | [0.560,1.212] | 9.14E-06 |
| Other Drugs SL | 0.949 | 1 | [0.622,1.448] | 0.0002 |
| ICB datasets, N = 152 (Figure 3d) | | | | |

| Biomarker | OR | p | 95% CI | p value under H0: ENLIGHT does not have a greater OR than the respective marker (one sided) |
|---|---|---|---|---|
| ENLIGHT | 2.69 | 0.037 | [1.24,4.59] | |
| gm(ENLIGHT,IFNG) | 4.076 | 0.002 | [1.946,8.539] | 1 |
| SELECT | 1.798 | 0.094 | [0.904,3.57] | 0.208 |
| ENLIGHT-InVitro | 0.832 | 1 | [0.409,1.689] | 0.01 |
| Target Expression | 2.111 | 0.088 | [1.06,4.204] | 0.312 |
| Other Drugs SL | 0.77 | 1 | [0.365,1.722] | 0.01 |
| Random Genes | 1.031 | 0.272 | [0.524,2.027] | 0.024 |
| Proliferation | 2.26 | 0.094 | [0.98,5.213] | 0.386 |
| Cytolytic | 2.342 | 0.092 | [1.041,5.270] | 0.406 |
| gm(IFNG,Cytolytic,Proliferation) | 2.039 | 0.088 | [4.018,6.459] | 0.285 |
| IFNG | 2.543 | 0.037 | [1.291,5.012] | 0.454 |
| Exhausted T-cell | 1.733 | 0.119 | [0.882,3.404] | 0.183 |
| TIDE | 1.144 | 0.271 | [0.586,2.234] | 0.038 |
| CD8+ T-cell | 1.622 | 0.139 | [0.831,3.167] | 0.147 |

**Table S4**. OR for ENLIGHT and other biomarkers. Each sub-table contains the OR, the p value for a test of the OR being greater than 1, the 95% CI of the OR and the p value for a test of greater OR for ENLIGHT vs. the corresponding biomarker (one sided).

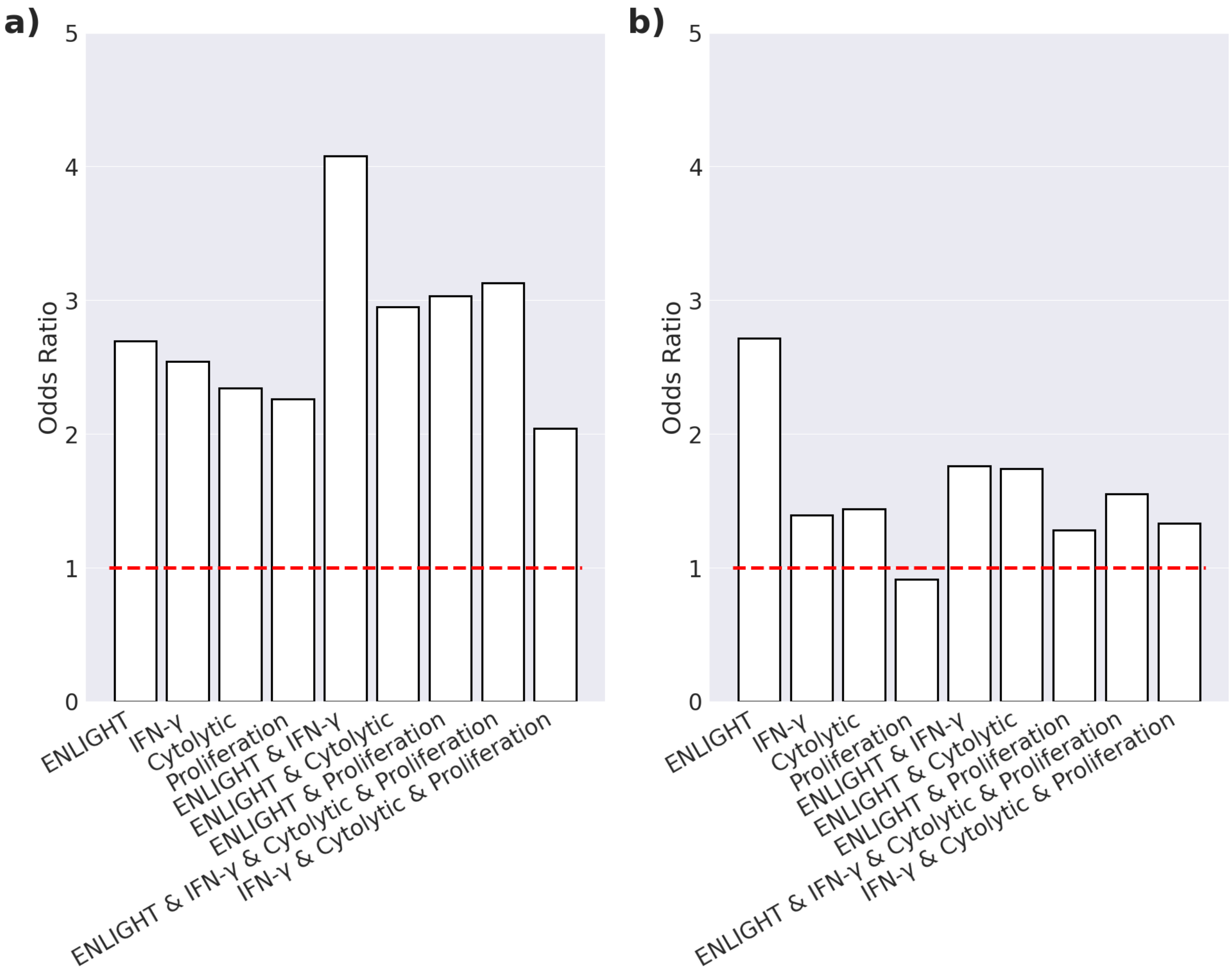


**Figure S4.** Comparison of OR between ENLIGHT and IFN-γ, Cytolytic and Proliferation signatures along with the combination between ENLIGHT and the three signatures on ICB datasets (**a**, N= 152) and non-ICB datasets (**b**, N = 511). X & Y refers to the geometric mean between the X and Y signatures per patient. For each signature, the decision threshold was calibrated as described in **STAR METHODS** (except *ENLIGHT* where the 0.54 threshold was used). The red dashed line represents an OR of 1 which is expected by chance.
